# SteMClass: A DNA Methylation-Based Classifier for iPSC In Vitro Differentiation States

**DOI:** 10.1101/2025.09.02.673063

**Authors:** Eilís Perez, Valeria Fernandez Vallone, Daniel Teichmann, Philipp Euskirchen, Christin Siewert, Brendan Osberg, Peggy Wolkenstein, Sebastian Diecke, Harald Stachelscheid, David Capper

## Abstract

Human induced pluripotent stem cells (iPSCs) hold great promise for regenerative medicine, disease modelling, and drug discovery, but most downstream applications require differentiation into specialised cell types not covered by current quality control assays. Here, we present “SteMClass”, a prototype DNA methylation-based classifier that standardises iPSC differentiation state identification across protocols with one test. We curated a reference cohort of 15 iPSC lines differentiated into seven distinct states (n = 97), performed array-based DNA methylation profiling, and trained a random forest model to classify the eight distinct differentiation states. In nested cross-validation, SteMClass achieved a Brier score of 0.018, and on an independent cohort (n = 58) attained 96.5% accuracy (Cohen’s K = 0.959) with a 3% rejection rate. Applied to external data (n = 249), SteMClass achieved 85.1% overall accuracy (Cohen’s K = 0.687) with a 12.9% rejection rate. Among classified samples (n = 217), accuracy was 97.7% (Cohen’s K = 0.93). SteMClass is compatible with all Illumina methylation array versions, and accessible via an interactive web interface that supports classification and exploration of DNA methylation profiles. By providing a harmonised, single-assay framework for iPSC-derived differentiation state characterisation, SteMClass improves reproducibility and comparability across studies, paving the way for robust quality control standards and accelerating clinical translation.

## Introduction

Due to their potential for self-renewal and to differentiate into all cell types of the human body, human induced pluripotent stem cells (iPSCs) hold immense promise for regenerative medicine, drug screening and disease modelling (Shi et al., 2017). Despite the potential of iPSCs, there are significant challenges to address before routine clinical use.

Key limitations include safety concerns regarding genomic stability, immunogenicity and the scalability and reproducibility of differentiation and characterisation methods. Many recent developments have been made in the field of iPSC production in order to combat these issues, such as the development of universal cells that are camouflaged from the immune system (Lanza et al., 2019), the use of small molecules to replace growth factors (Liuyang et al., 2023; Wang et al., 2024) and improved quality control of iPSC production such as genetic screening, and potency assessment (Dobner et al., 2024; Lenz, 2012, 2012; MacArthur et al., 2019; Tsankov et al., 2015; Yoshihara et al., 2017, Kügelgen et al., 2025). Nevertheless, scalable and reproducible differentiation of iPSCs into standardised, well-defined cell types remains a key challenge.

In recent years, large cell and tissue atlases have been generated using gene expression and DNA methylation profiling to identify distinct cell populations in vivo (Loyfer et al., 2023; Moss et al., 2018). However, cells generated by the differentiation of iPSCs in vitro are inherently distinct from their terminally differentiated in vivo counterparts due to their developmentally immature state, in vitro cell culture associated changes, reprogramming artefacts, and the persistence of somatic cell-specific epigenetic signatures (Cerneckis et al., 2024; Diane et al., 2025; Edgar et al., 2022; Franzen et al., 2021). Conventional assessments of iPSC differentiation generally depend on measuring a limited panel of lineage-characteristic gene or protein markers. Because marker choice and interpretation vary, this strategy is susceptible to considerable variability across laboratories (Grancharova et al., 2021; Kramer et al., 2016). Using long-read nanopore transcriptomics of trilineage-differentiated samples from 15 iPSC lines, Dobner et al. showed that even ISSCR-endorsed markers overlap significantly across germ layers and developed a 12-gene machine-learning classifier for early lineage identity (Dobner et al., 2024; Ludwig et al., 2023).

DNA methylation is tightly regulated and varies according to cell type with dynamic alterations playing a critical role in guiding cell fate and development. Established during development and maintained through cell division, DNA methylation serves as a more consistent and enduring marker of a cell’s identity than gene expression, which can be transient and subject to rapid changes in response to external stimuli (Bird, 2002; Gibney & Nolan, 2010; Jones & Taylor, 1980; Tompkins, 2022). DNMT-mediated methylation and TET-mediated turnover are fundamental as cells exit pluripotency (Parry et al., 2021). Research indicates that the efficiency of iPSC differentiation and the stability of differentiated cell populations are influenced by DNA methylation (Buckberry et al., 2023; Hajmousa et al., 2024; Ideno et al., 2022; Tesarova et al., 2016; Yang et al., 2023) and that DNA methylation patterns can serve as markers to identify iPSC lines with varying differentiation capacities (Butcher et al., 2016; Choi et al., 2020; Edgar et al., 2022; Nishizawa et al., 2016; Schmidt et al., 2022). Therefore, understanding the dynamics of DNA methylation changes through differentiation is key to improving differentiation efficiency and the standardisation of producing iPSCs and their differentiation derivatives.

While whole-genome bisulfite sequencing (WGBS) provides a comprehensive overview of the methylome, array-based methods such as the Illumina Infinium assay, offer a cost-effective alternative for large-scale studies, enabling quantitative analysis of hundreds of thousands of curated CpG sites with high reproducibility and minimal inter-user variability (Bibikova et al., 2006; Capper et al., 2018; Chirica et al., 2024; Kurdyukov & Bullock, 2016; Tran et al., 2024). Combined with machine learning, the scalability of methylation arrays has enabled robust classifiers for several tumour types (Capper et al., 2018; Danielsson et al., 2015; Dragomir et al., 2023; Hoang et al., 2024; Jurmeister et al., 2022; Tran et al., 2024). In brain tumour diagnostics, harmonised methods, reference datasets and interpretive criteria made methylation data comparable across laboratories and the resulting classifier transformed clinical practice and was formally adopted by the WHO (Louis et al., 2021).

In this work, we bring these concepts to iPSC research. Using a centrally curated, publicly available reference dataset, we trained a random forest classifier, SteMClass, that identifies defined stages of iPSC differentiation from DNA methylation profiles. Researchers can upload their own test data via an online interface (https://stemclass.charite.de/) and assess differentiation state against the reference with a single assay, rather than multiple targeted tests. By providing a common benchmark across studies and protocols, SteMClass could promote harmonisation in iPSC research and improve the reproducibility and interpretability of findings, a key requirement for the quality control of iPSC-derived cells intended for clinical use.

## Materials and Methods

### Cell Culture and Differentiation experiments

#### Culture of iPSCs

Human induced pluripotent stem cells (iPSCs) were maintained on growth factor reduced Geltrex (Gibco) in E8 or mTeSR (Stem Cell Technologies) media with daily media exchange. Cells were EDTA clump-passaged every 2-4 days as previously described (Fernandez Vallone et al., 2025). iPSC lines, culture conditions, and outcome of G-banding and CNV analysis (Kügelgen et al., 2025) are listed in Supplementary Table 1, individual karyotyping results are available upon request. iPSC lines used for StemClass development were provided by the Core Unit Pluripotent Stem Cells and Organoids of the Berlin Institute of Health at Charité (CUSCO) and were generated under ethical approval following informed consent from the donors.

#### Differentiation Protocols

Cell type nomenclatures, iPSC lines and corresponding differentiation protocols used in this study are described in Supplementary Tables 1-3 and Supplementary Figure 1.

iPSCs were differentiated to ectoderm, neural stem cells (dual SMAD inhibition) and astrocytes (astrocytes adapted from TCW et al., 2017, with GLAST-positive selection), definitive endoderm (STEMdiff™ Definitive Endoderm Kit), lung progenitors (adapted from Jacob et al., 2019, with CPM-positive selection), mesoderm and endothelial cells (adapted from Patsch et al., 2015, with CD144/CD31 selection), and by trilineage differentiation (StemMACS™ Trilineage Differentiation Kit). Differentiation efficiency was assessed by flow cytometry for lineage markers, and only populations meeting predefined purity thresholds (>80-90% marker-positive) were used. Full protocols, reagents and flow cytometry procedures are provided in the Supplementary Experimental Procedures, cell type nomenclatures, iPSC lines and protocols are summarised in Supplementary Tables 1-3 and Supplementary Figure 1.

### DNA Extraction

DNA was extracted from cell pellets using the Maxwell® RSC Instrument with the Cultured Cells, Blood or FFPE Plus DNA kits (Promega) as appropriate to sample type, with no difference in methylation data quality, and quantified with the QuantiFluor® ONE dsDNA kit (Promega).

### DNA Methylation Analysis

A cohort of n=155 iPSCs and iPSC derivates of seven differentiation states was generated in technical (independent experiments) or biological (distinct cell line donors) replicates, then randomly split into a training cohort of n=97 with a minimum of 8 samples per differentiation state, and a validation cohort of n=58. DNA extraction was performed on each sample cell pellet for subsequent DNA methylation profiling.

### Array-Based DNA Methylation Profiling

Genomic DNA was bisulfite-converted using the EpiTect Fast Bisulfite Kit (Qiagen) following the manufacturers’ protocol. The bisulfite-converted DNA was subsequently analysed using the Infinium MethylationEPIC BeadChip array (v1.0 or v2.0) and scanned on the iScan System (Illumina). Signal intensities were obtained from IDAT files using the Bioconductor package minfi (v 1.40.0) (Aryee et al., 2014). Dye bias correction was performed using the Noob (normal-exponential out-of-band) method with the preprocessNoob function without “between-array normalisation” (Triche et al., 2013). Sex prediction of samples was performed using the predictSex function of the wateRmelon package (v 1.36.0). Copy-number variation analysis from methylation array data was performed using the conumee 2.0 Bioconductor package (v.2.1.2) (Daenekas et al., 2024). A cohort of 50 control samples displaying a balanced copy-number profile from female donors were used for normalization, to allow for inference of sex from the copy number of the X chromosome.

### Analysis of Publicly Available Datasets

A publicly available methylation array dataset was systematically curated and downloaded from Gene Expression Omnibus (GEO) as raw IDAT files (accessions listed in Supplementary Table 5). The PRISMA flow chart of dataset inclusion and exclusion shown in Supplementary Figure 7 was performed according to Page et al. (Page et al., 2021). GenomicMethylSet objects were generated from raw signal intensity files using the readGEORawFile function of the GEOquery package (v2.62.1) (Davis & Meltzer, 2007). Data filtering steps included the removal of probes with detection p-values greater than 0.05 in more than 10% of samples, probes containing single nucleotide polymorphisms (SNPs) at the CpG site, probes located on sex chromosomes and cross-reactive probes as previously described (Capper et al., 2018, Chen et al., 2013). Raw data from EPIC arrays were merged with publicly available 450K array datasets using the combineArrays function in minfi. The combined dataset underwent the same filtering and normalisation process as described above.

### Differential Methylation Analysis

Differential methylation analysis on the CpG site and region level was performed in a one-to-one fashion between iPSCs and the respective differentiation states as previously described (Maksimovic et al., 2017). Methylation-specific gene ontology (GO) analysis of differentially methylated positions (DMPs) was conducted for each differentiation state using the gometh function within the missMethyl package (v 1.28.0) (Phipson et al., 2016). “Broader profiles” beyond the classifier features were defined as the top 20,000 CpGs from our DMP analysis that distinguished each reference differentiation state from iPSC, for iPSC it was the top 20,000 CpGs that were common to all differentiation state vs iPSC comparisons. The annotated tables for the top 20,000 CpGs per differentiation state are available in Supplementary Table 4, full DMP analysis results tables are available on request.

Temporal methylation modules were identified separately for astrocyte and lung differentiation using limma (v. 3.62.2) linear models on M-values, with time point as a categorical variable and parental line as a covariate. CpGs with significant temporal variation (moderated F-test, Benjamini-Hochberg-adjusted P < 0.05) and a range of ≥0.10 in mean beta value across stages were Z-scored across stages and grouped by k-means clustering (100 random starts) into decreasing (M1) and increasing (M2) modules. Full module tables are available on request.

### Unsupervised Analysis

Principal component analysis (PCA) was performed on DNA methylation beta values. Separate analyses were conducted for 4 time points throughout astrocyte differentiation across three parental iPSC lines (n = 12), 6 time points throughout lung differentiation across two parental iPSC lines (n = 12). An additional PCA was performed for 9 time points throughout one astrocyte differentiation with cohorts of reference iPSC and astrocyte samples (n = 35). For each analysis, CpGs with missing or non-finite values were excluded, and the 5,000 most variable CpGs were selected based on their standard deviation across samples. Consecutive stage centroids were connected chronologically to illustrate differentiation trajectories.

Uniform Manifold Approximation and Projection (UMAP) was performed on the top 50,000 most variable CpGs for each respective dataset using the R package umap (v 0.2.10.0). UMAP projections were computed using the uwot R package (v 0.2.2). UMAP parameters were set to n_neighbors=18, min_dist=0.3 n_components=2, with a Euclidean distance metric and a fixed random state for reproducibility.

### Classifier development

A random forest classifier was built with ranger (v 0.16.0) within the tidymodels ecosystem (v 1.2.0) under R 4.3.3 (further package versions in the Supplementary Experimental Procedures). A nested cross-validation framework was used to obtain unbiased estimates of model performance while preventing information leakage (5-fold outer cross-validation repeated three times, 3-fold inner cross-validation).

#### Nested cross validation, feature selection and hyperparameter tuning

Within each outer training split, CpG sites were first variance-filtered using the training data only, retaining the top 50,000 CpGs ranked by standard deviation to reduce dimensionality. Random forest hyperparameters were tuned exclusively in the inner cross-validation using a 20-point max-entropy (Latin hypercube) design over minimum node size (2-30) and number of trees (300-2000), while mtry was fixed per split to the square root of the number of retained predictors. Hyperparameters were selected by minimising the multiclass Brier score. The model was then refit on the outer training split, CpGs were ranked by permutation importance, and nested feature selection fitted models on increasing prefixes of this list (2,000, 5,000, and 10,000 CpGs) using the same inner cross-validation structure. The final feature count per outer split was determined using a one-standard-error plateau rule on the inner-CV Brier score. Across outer splits, selected CpGs were aggregated by selection frequency. The final deployed feature set comprised the top-ranked CpGs up to the median selected feature count across splits. Consensus hyperparameters derived from the nested procedure were then used to refit the model on all training data.

#### Probability score calibration and threshold identification

For probability calibration, outer fold predictions were pooled, and a multinomial ridge regression model was trained in a cross-fitted manner: predictions for each outer fold were calibrated using a model trained only on predictions from the remaining outer folds. Calibration performance was summarised using the multiclass Brier score. The rejection threshold was selected as the largest probability cut-off on the macro-averaged Youden index plateau of the pooled cross-fitted calibrated predictions, i.e. before a statistically significant decrease (95% bootstrap confidence interval), samples below it were labelled “Not Classifiable”.

#### Final model and variable importance

The final random forest model was trained on the entire training cohort using the final feature set (10,000 CpGs) and consensus hyperparameters (mtry= 100, min_n = 5, trees = 1192) with a calibrated probability score threshold of 0.5. Variable importance was extracted from the fitted ranger object using the impurity-based measure.

#### Model evaluation

Performance was assessed on the internal (n = 58) and external (n = 249, publicly available array data) validation cohorts (Supplementary Table 5). For validation samples with discordant classifications, the CpG features contributing most negatively toward the annotated class and most positively toward the SteMClass class were extracted from per-sample feature contributions computed on the tidy forest object.

#### SteMClass Web Application

The SteMClass web application was developed in R Shiny. Uploaded IDAT files are noob-normalised, harmonised to the 450K format where required, aligned to the training reference with mean imputation of missing CpGs and classified by the pre-trained model. Outputs include class probabilities with a rejection option, UMAP projections and global and gene-level heatmaps (implementation details in the Supplementary Experimental Procedures).

#### Cumulative-contribution analysis of CpG features

Per-sample feature contributions were computed with featureContrib from tree.interpreter (v 0.1.1) and averaged as mean absolute contributions per CpG and class. For each class, CpGs were ranked by this value and the cumulative fraction of total contribution was plotted against CpG rank (log10 scale, details in the Supplementary Experimental Procedures).

### Nanopore Sequencing

Data set used comprised an internal cohort (n= 16 iPSC lines plus trilineage derivatives) and an external cohort of samples derived from trilineage differentiation of n= 12 different iPSC lines. External cohort was kindly provided by Dr. Sebastian Diecke (Max Delbrück Center, MDC).

Libraries were prepared with the Rapid Barcoding Kit (SQK-RBK004 or SQK-RBK114.24; Oxford Nanopore Technologies) and sequenced at low coverage on R9.4.1 or R10.4.1 flow cells on a MinION Mk 1B, multiplexing up to 6 samples per run. Analysis used the nanoDx pipeline (v0.6.2, Kuschel et al., 2022, https://gitlab.com/pesk/nanoDx) with the SteMClass training cohort converted to H5 format and copy number profiling used ichorCNA (v.0.5.1) (Adalsteinsson et al., 2017). Full library preparation and run details are provided in the Supplementary Experimental Procedures.

#### Nanopore sequencing data processing

Raw signal was basecalled with dorado (high-accuracy 5mCG_5hmCG model) and aligned to hg19 with minimap2, per-CpG methylation was summarised with modkit pileup, combining 5mC and 5hmC to match bisulfite-based arrays (software versions and filtering in the Supplementary Experimental Procedures).

#### Classification and score stability analysis

Samples were classified with the nanoDx pipeline (v0.6.2, Euskirchen et al., 2017) using an ad hoc random forest trained, per sample, on the SteMClass reference set (97 samples, 8 classes, 10,000 CpGs) restricted to the CpGs covered by that sample, with sigmoid calibration. The 0.5 confidence threshold was prespecified from the array classifier, samples below it were designated Not Classifiable. To determine how much sequencing data a stable SteMClass classification requires, each sequencing run from the validation cohorts which resulted in a classification score meeting the 0.5 cut off was reanalysed retrospectively with increasing amounts of sequencing data (n =63) (Supplementary Experimental Procedures).

## Results

### DNA methylation remodelling accompanies lineage-specific differentiation of iPSCs in vitro

DNA methylation is a well-established feature of cell- and tissue-type identity *in vivo*. To investigate whether differentiation-associated methylation patterns are similarly established *in vitro*, we examined whether genetically identical cells could be distinguished by their differentiation state and whether these changes were consistent across independent parental iPSC lines. We analysed DNA methylation profiles from two iPSC lines differentiated towards lung progenitors and three iPSC lines differentiated towards an astrocytic lineage (Supplementary Figure 1a & c).

Principal component analysis (PCA) of the 5,000 most variable CpG sites revealed that DNA methylation profiles were strongly associated with differentiation stage in both the astrocyte and lung differentiations (Figure 1). For astrocytes, samples collected at the iPSC stage and day 7 clustered closer together, whereas days 19 and 37 formed distinct groups along PC1, indicating substantial methylation remodelling between days 7 and 19 (Figure 1a). Lung differentiation similarly showed stage-associated progression along PC1, with samples ordered from day -2 (iPSC) through day 15 (Figure 1b). In both differentiations, the stage-associated trajectory along PC1 was consistent across parental lines. PC2 instead captured differences between iPSC lines. In the lung differentiation, PC2 completely separated BIHi001-B from BIHi005-A at every time point, from the iPSC stage onwards. Day -2 and day 3 samples from both lines were processed on the same array slide yet still separated just as clearly on PC2, therefore the difference must come from the cell lines themselves rather than from technical variation between slides. Line-related variation accounted for less of the methylation variance in the astrocyte differentiation (PC2: 2.6%) than in the lung differentiation (PC2: 8.0%). As variance explained is relative, this difference likely reflects larger differentiation-associated methylation changes during astrocyte differentiation, which would reduce the share of the 5,000 most variable CpGs, and of the total variance, attributable to line effects.

**Figure 1:**
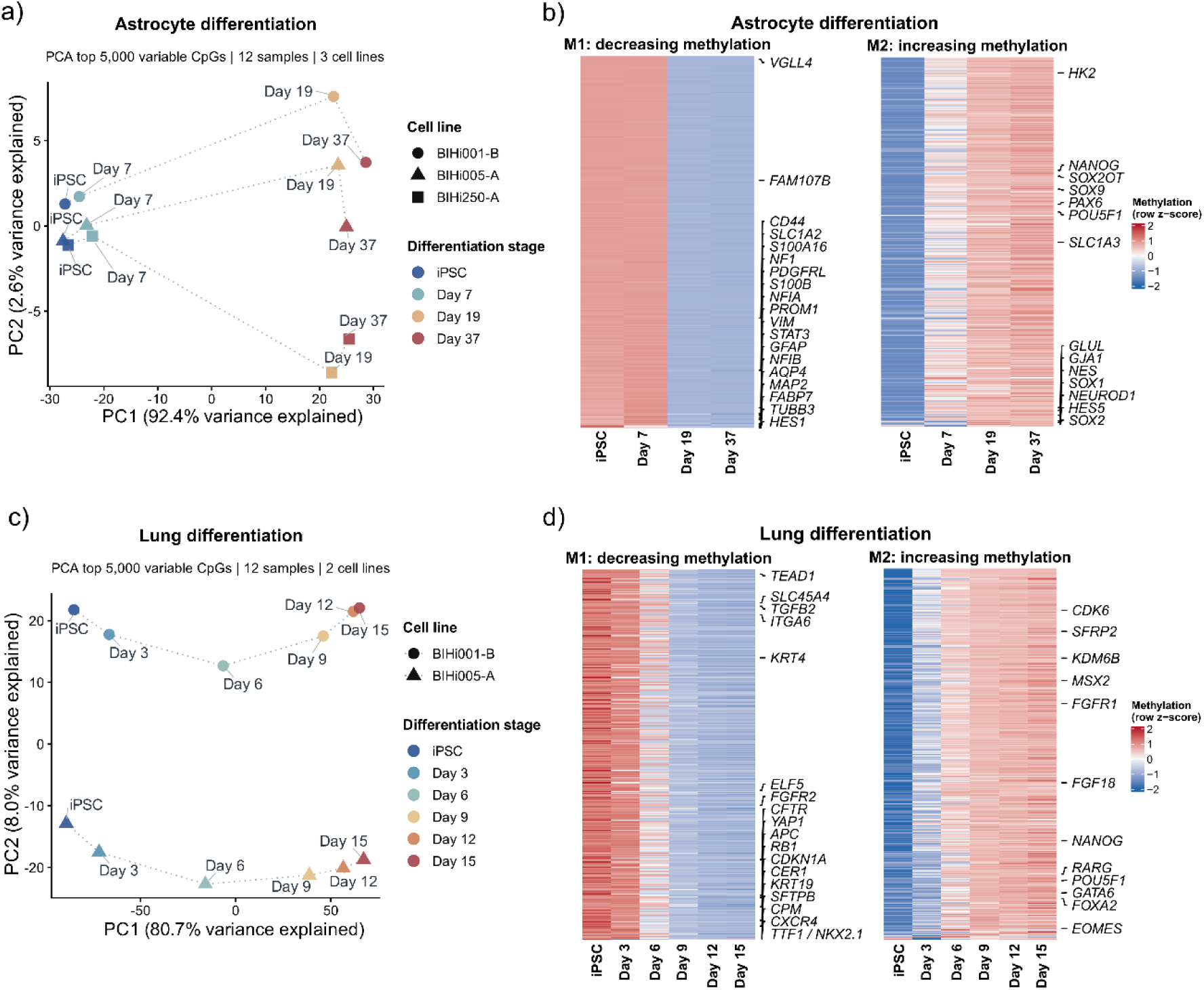
Identification of progressive DNA methylation changes during in vitro iPSC differentiation. (a) Principal component analysis (PCA) of DNA methylation profiles during astrocyte differentiation across four timepoints repeated in three iPSC lines, based on the 5,000 most variable CpG sites. Samples are coloured by differentiation stage and distinguished by iPSC line using point shapes. PC1 and PC2 explain 92.4% and 2.6% of the total variance, respectively. (b) Heatmaps showing two temporal DNA methylation modules during astrocyte differentiation. M1 and M2 comprise CpGs with decreasing and increasing methylation, respectively. Columns represent differentiation stages (iPSC, day 7, day 19 and day 37), with methylation values averaged across samples at each stage. Rows represent individual CpG sites, and colours indicate row-wise Z-scored mean methylation β-values. Selected CpGs are labelled with their associated genes. (c) PCA of DNA methylation profiles during lung progenitor differentiation across four timepoints repeated in two iPSC lines, displayed as in a). PC1 and PC2 explain 80.7% and 8.0% of the total variance, respectively. (d) Temporal DNA methylation modules during lung progenitor differentiation, displayed as in (b). Columns represent differentiation stages (iPSC, day 3, day 6, day 9, day 12 and day 15).

To characterise the underlying CpG-level changes, we identified two temporal methylation modules in each differentiation system, representing coordinated losses (M1) and gains (M2) of methylation (Figure 1c & 1d). During astrocyte differentiation, M1 included CpGs annotated to astrocyte-associated markers, including *GFAP*, *S100B*, *S100A16*, *CD44* and *SLC1A2*, whereas M2 contained sites associated with the pluripotency markers *POU5F1* and *NANOG*, alongside *HK2*, *SOX2OT* and the neuroectodermal marker *PAX6*. The most pronounced methylation changes occurred between days 7 and 19, in agreement with the PCA (Figure 1a & c). During lung differentiation, M1 included CpGs annotated to lung-associated markers (*KRT19*, *CFTR*, *FGFR2*, *CPM* and *NKX2.1*) and the definitive endoderm marker *CXCR4*, whereas M2 contained sites associated with *POU5F1*, *NANOG* and the developmental regulators *GATA6*, *EOMES* and *FOXA2* (Figure 1d). Flow cytometry additionally demonstrated increasing expression of CXCR4, NKX2.1 and CPM over the course of lung differentiation (Supplementary Figure 2a).

To examine the temporal progression of methylation remodelling in greater detail, we additionally analysed a more densely sampled astrocyte differentiation experiment comprising nine time points. PCA incorporating a reference cohort of iPSCs (n = 14) and their differentiated astrocyte derivatives (n = 12) revealed clear separation between the endpoint states, with intermediate samples distributed along the differentiation axis (Supplementary Figure 2b). Genome-wide comparison of iPSCs and differentiated astrocytes identified 228,765 differentially methylated positions (DMPs, Supplementary Table 4). Promoter-associated CpGs annotated to astrocyte-associated genes, including *GFAP*, *S100B* and *S100A16*, exhibited reduced methylation in differentiated astrocytes, whereas promoter-associated CpGs linked to *POU5F1* and *HK2* gained methylation (Supplementary Figure 2c). Examination of these loci across the nine-point time course revealed that methylation changes occurred at distinct stages. CpGs associated with *POU5F1* and *HK2* began gaining methylation by days 3-5, preceding the loss of methylation at *S100A16*- and *S100B*-associated CpGs from approximately day 14 onwards. Demethylation at the *GFAP* locus occurred later in differentiation (Supplementary Figure 2c). Notably, a gene-body CpG annotated to PAX6 (cg02770983), a canonical neuroectodermal marker gene, was also among the differentiation-associated DMPs and gained methylation by days 3-5 (Supplementary Figure 2c).

Collectively, these findings demonstrate that differentiation of isogenic iPSCs is accompanied by coordinated, stage-associated DNA methylation remodelling that is consistent across independent parental lines and lineages.

### Unsupervised dimensionality reduction of DNA methylation data separates iPSC-derived cells according to differentiation state

We next investigated whether the lineage-specific methylation changes observed during astrocyte and lung differentiation generalise to other iPSC derived cell types. Early ectodermal, mesodermal and endodermal progenitor cultures were generated with at least two independent protocols per lineage from 11 independent iPSC lines (see Methods). To minimise heterogeneity, only populations with ≥85% marker-positive cells by FACS were analysed (Supplementary Figures 1 and 3, Supplementary Table 2). Unsupervised analysis of the 50,000 most variable CpG sites across all samples was visualised with UMAP (Figure 2a). Four discrete clusters emerged: pluripotent iPSCs and three well-separated groups corresponding to the three germ-layers ectoderm, mesoderm and endoderm. Samples from different protocols converged on the same germ-layer cluster and replicates from separate batches co-localised, demonstrating robust, reproducible signatures.

**Figure 2:**
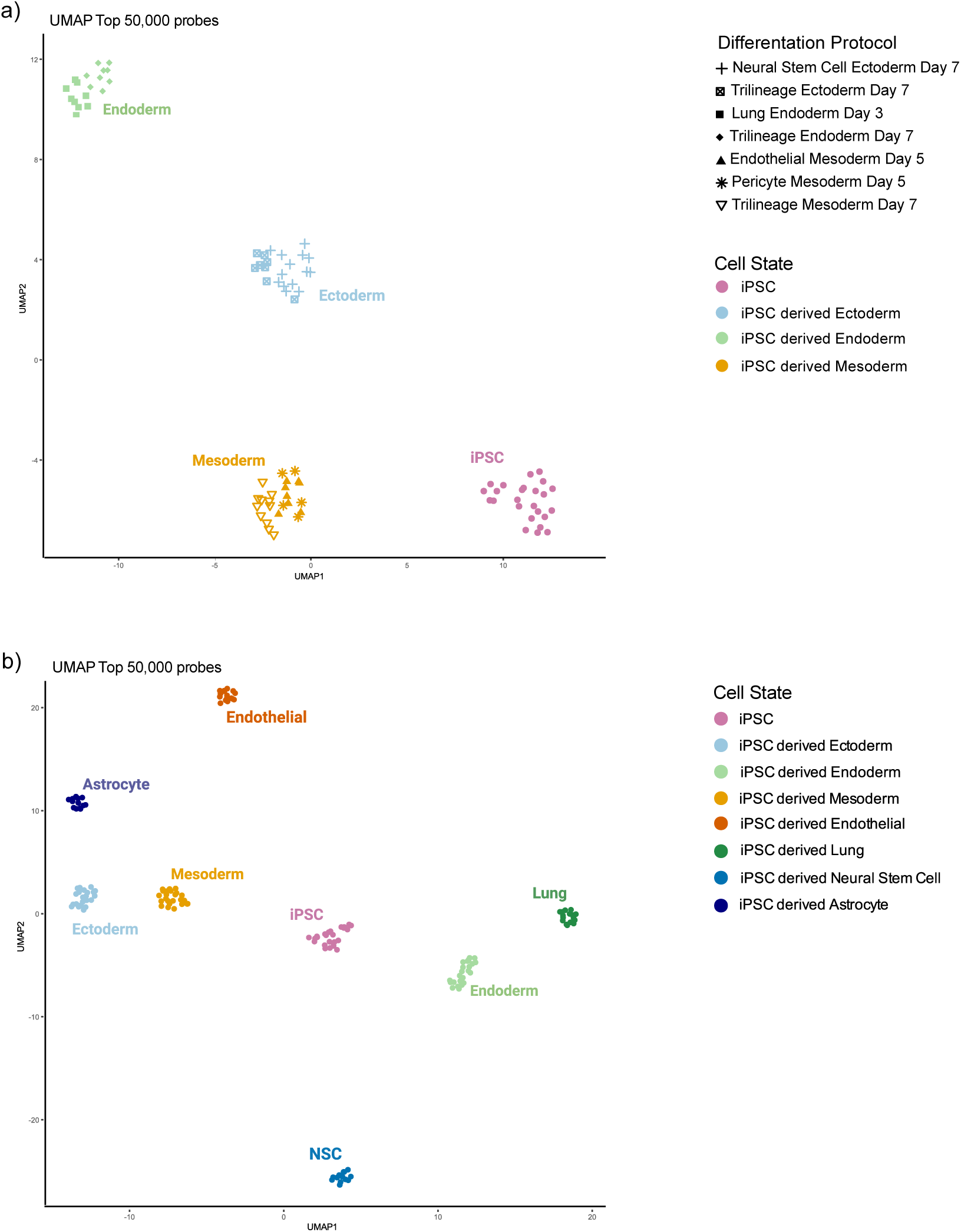
Unsupervised analysis indicates that differentiation state-specific DNA methylation signatures are reproducible across cell lines and protocols. **(a)** UMAP plot of DNA methylation data (50,000 most variable CpGs) from 15 independent iPSC lines, 11 of which were differentiated into ectodermal, endodermal, and mesodermal lineages using at least two differentiation protocols per lineage with each point representing an independent technical and/or biological replicate. Colours represent differentiation states, the differentiation protocol used is indicated by shape. **(b)** UMAP plot as in (a), including at least one additional differentiation state per germ layer. Colours represent differentiation states; iPSCs are represented in pink, endodermal differentiation states (endoderm and lung) are represented in green tones, ectodermal differentiation states (ectoderm, neural stem cells and astrocytes) are represented in blue tones and mesodermal differentiation states (mesoderm and endothelial cells) are represented in orange tones.

**Figure 3.**
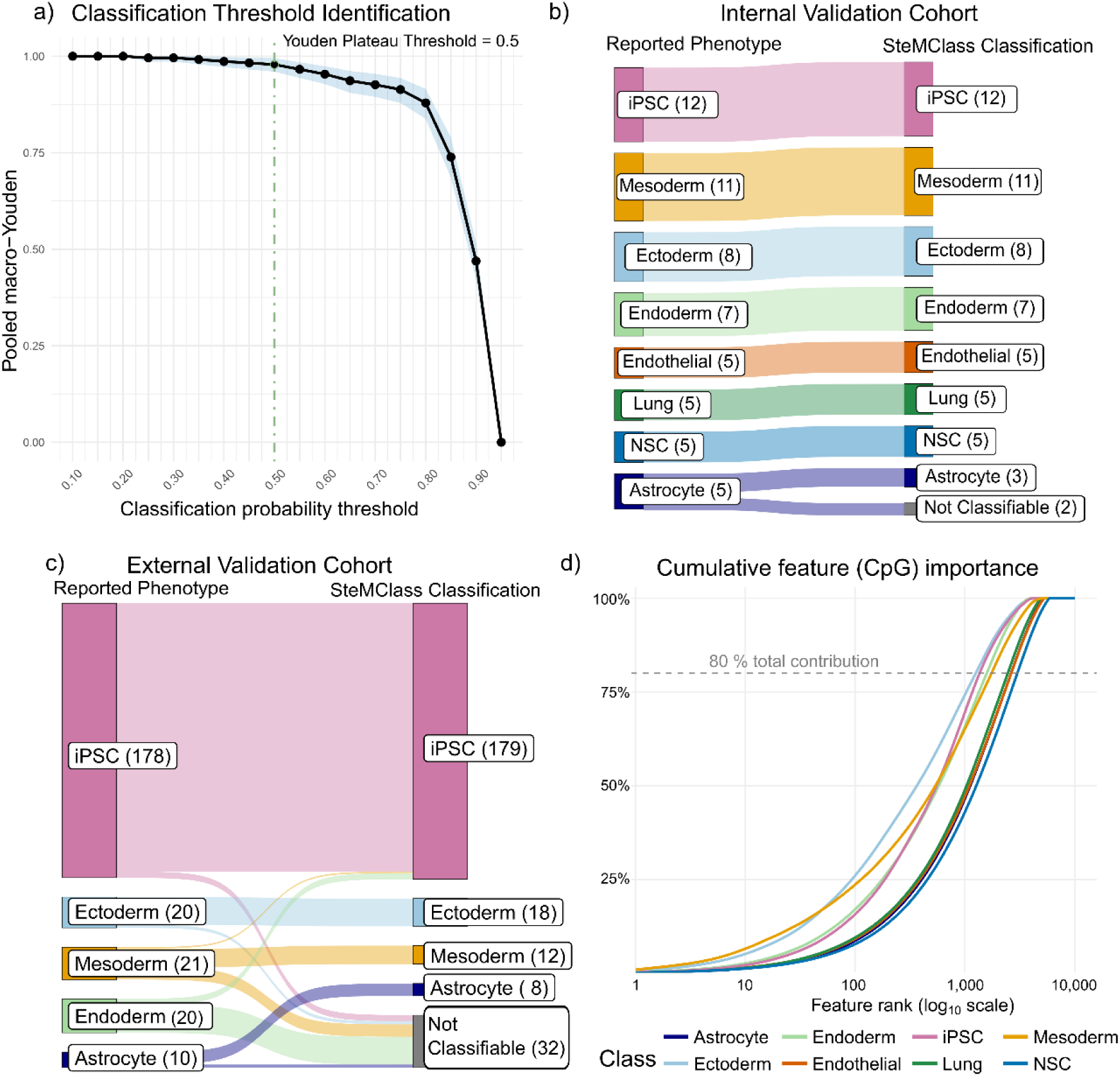
Integrated assessment of SteMClass: threshold tuning, independent validation, and feature-level signal distribution. **(a)** Macro-Youden index as a function of classification probability threshold. Points denote empirical estimates computed from pooled cross-validated calibrated predictions and the ribbon shows the 95% percentile bootstrap confidence interval (1,000 resamples). The vertical dashed line indicates the largest threshold before a statistically significant improvement when lowering the cut-off. Significance was assessed using paired bootstrap differences between adjacent thresholds (one-sided test on ΔYouden). **(b)** Sankey diagram showing classifier performance on the independent internal validation cohort (n = 58). Each flow connects the original sample annotation (reported phenotype) to the SteMClass predicted class. The internal cohort comprised: Astrocytes (n = 5), Ectodermal cells (n = 8), Endodermal cells (n = 7), Mesodermal cells (n = 11), Endothelial cells (n = 5), iPSCs (n = 12), Lung cells (n = 5), and NSCs (n = 5). **(c)** As in (b), for the external validation cohort. The external cohort comprises iPSCs (n = 178), iPSC-derived Ectoderm (n = 20), Endoderm (n = 20), Mesoderm (n = 21) and Astrocytes (n=10) from multiple studies (Supplementary Table 5). **(d)** Cumulative absolute feature contribution per class. Lines show the summed absolute contribution per class as CpGs are added in descending order of mean absolute contribution across all 307 test samples. The x-axis is on a log₁₀ scale, the dashed horizontal line marks 80% of the total contribution.

To determine whether DNA methylation signatures can resolve a broader spectrum of differentiation states, we expanded the dataset to a total of eight in vitro cell states (undifferentiated iPSCs, ectodermal cells, mesodermal cells, endodermal cells, lung cells, endothelial cells, neural stem cells, and astrocytes). Representative marker expression and morphology data are shown in Supplementary Figure 3.

Unsupervised analysis of the 50,000 most variable CpG sites across all samples, visualised with UMAP, produced eight well-separated clusters that mapped one-to-one onto the designated cell states (Figure 2b). The observed clustering of distinct differentiation states was independent of potential batch effects such as parental cell line, donor sex, or DNA methylation array (EPIC) version (Supplementary Figure 4a-d). Pairwise differential methylation analyses of iPSCs against each of the differentiation states, combined with gene ontology enrichment, confirmed that the methylation changes align with known lineage-specific biological processes in each differentiation state (Supplementary Figure 5 and 6).

These findings demonstrate that genome-wide DNA methylation profiles provide a robust read-out of differentiation state across a diverse set of iPSC lines and differentiation derivatives, underscoring their utility for benchmarking and standardising in vitro differentiation characterisation.

### Building and Validating a DNA Methylation-Based Differentiation State Classifier: SteMClass

On observing that the methylation differences between in vitro differentiation states were reflective of differentiation processes, we leveraged this to train a random forest algorithm for DNA methylation-based differentiation state classification. Random forest models suit the correlated, high-dimensional structure of methylation data through implicit handling of collinearity and built-in feature selection. Model development used nested cross-validation for unbiased performance estimation, feature selection, hyperparameter tuning, calibration and threshold determination (Methods, Supplementary Figure 7a). Cross-fitted calibrated probabilities obtained using a multinomial ridge regression calibrator yielded a multiclass Brier score of 0.018 and a macro expected calibration error (ECE) of 0.028, indicating well-calibrated predictions (Supplementary Figure 7b). To reduce overconfident classifications, we evaluated candidate rejection thresholds between 0.10 and 0.95 using the pooled cross-fitted calibrated predictions and the macro-averaged Youden index. The selected operating point was a probability cut-off of 0.5 (macro-Youden = 0.978), corresponding to the highest threshold before a statistically significant decline in macro-Youden was observed (Figure 3a). Lowering the threshold from 0.55 to 0.50 significantly improved Macro-Youden (p = 0.00899), whereas lowering from 0.50 to 0.45 did not (p = 0.372). Samples with a maximum predicted class probability below 0.5 were therefore rejected and labelled “Not Classifiable”.

SteMClass was evaluated on an independent cohort of 58 samples representing the eight distinct differentiation states included in the classifier and attained 96.5% accuracy, Cohen’s K = 0.959, and a 3% reject rate (Figure 3b). These samples were generated in-house from the same iPSC lines and protocols as the training cohort, but represented independent experiments, reagent batches, and personnel.

To further evaluate SteMClass, we assembled an external validation cohort of 249 publicly available DNA methylation array samples (Supplementary Figure 8a). Sample labels were assigned based on the materials and methods described in the originating publications and mapped to the differentiation states defined in this study. These annotations derive from diverse protocols and levels of characterisation and therefore do not constitute a uniform ground truth. This heterogeneity reflects the current landscape of in vitro differentiation studies and represents precisely the scenario SteMClass was designed to address, objective, methylation-based state assignment across experimentally variable systems. In this context, relative to literature-reported annotations, SteMClass demonstrated 85.1% agreement (Cohen’s K = 0.687) with a 12.9% rejection rate (Figure 3c). Among classified samples (n = 217), accuracy was 97.7% (Cohen’s K = 0.93). To visualise how many CpG probes drive the classifier decisions, we quantified the absolute feature contribution of every probe for every test sample (internal and external combined) and aggregated these values across samples (see Methods). The resulting cumulative-contribution curves (Figure 3d) rise gradually: no class reaches 80% of its total weight before 1000 ranked probes and the remaining 20% is dispersed over several thousand additional sites. Hence SteMClass recognises each lineage through a wide methylation profile rather than a handful of high-impact markers, making the model inherently robust to noise at individual CpGs.

We also tested iPSC-derived samples from the same datasets annotated with differentiation states outside SteMClass’s training scope (n = 54; Supplementary Figure 8b, Supplementary Table 5). As expected for states outside the scope of SteMClass, most of these samples were not classified. Six early-stage endoderm (< 3 days) and three embryoid body samples were classified as iPSC. One embryoid body sample was classified as ectoderm and one lung progenitor sample (at a differentiation stage not represented in SteMClass) was classified as mesoderm, the remaining samples were not classifiable. The embryoid body assignments to iPSC or ectoderm occurred in developmentally proximal states and are therefore consistent with projection onto the closest reference lineage. As closed-set classifiers, multiclass random forests may assign out-of-distribution samples near decision boundaries to a neighbouring lineage. Across 303 external validation samples, only a single biologically discordant assignment (lung progenitor classified as mesoderm) was observed.

### SteMClass classification can be performed using Nanopore low pass whole genome sequencing

To assess the feasibility of SteMClass classification using Nanopore sequencing, 13 samples from the internal validation cohort (n = 58) were analysed using both EPIC methylation arrays and Nanopore sequencing (Supplementary Figure 8c). Nanopore-derived methylation profiles were classified using the nanoDx pipeline (Euskirchen et al., 2017), employing an ad hoc random forest (RF) model trained on the SteMClass reference set. Classification was concordant between platforms in 12/13 samples (92.3%). We additionally analysed 30 internal validation samples and 36 samples cultured, differentiated and DNA-extracted at an independent external site, using lineage annotations established by gold-standard markers as the reference (Supplementary Table 2). Applying the threshold of ≥0.5, 28/30 (93.3%) internal and 35/36 (97.2%) external validation samples were classifiable (overall 63/66, 95.5%), with 100% concordance with the lineage annotations among classifiable samples (Figure 4a & 4b). The three not classifiable samples had below-median sequencing depth and CpG coverage (77,000-125,000 reads and 282-515 CpGs, compared with cohort medians of 153,000 reads and 783 CpGs) (Supplementary Table 2).

**Figure 4.**
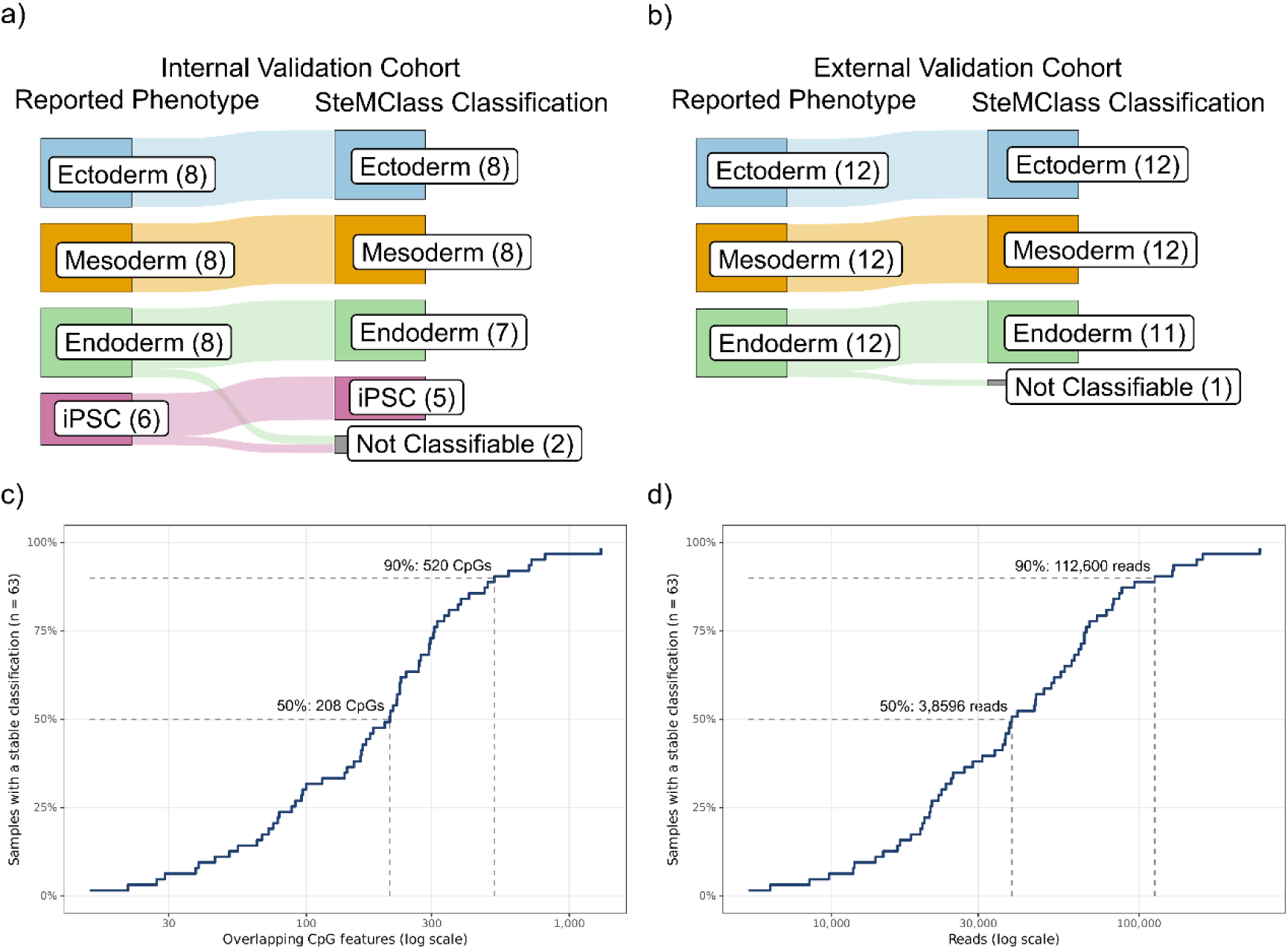
SteMClass classification of nanopore methylation profiles in independent validation cohorts and assessment of the sequencing data required for a stable classification in this cohort. **(a)** Sankey diagram showing classifier performance on the independent internal validation cohort (n = 30). Each flow connects the original sample annotation (reported phenotype) to the SteMClass predicted class. The internal cohort comprised ectodermal cells (n = 8), endodermal cells (n = 8), mesodermal cells (n = 9) and iPSCs (n = 6).(b) As in (a), for the external validation cohort (n = 36), cultured, differentiated and DNA-extracted at an external site. The external cohort comprised ectodermal cells (n = 12), endodermal cells (n = 12) and mesodermal cells (n = 12). **(c, d)** Cumulative proportion of the 63 classifiable samples that had reached a stable classification, plotted against (c) the number of overlapping CpG features and (d) the number of reads (log scale). Each step marks one sample reaching classification score stability, the curve’s height gives the proportion of samples stable at that amount of data. Dashed lines mark the depths at which 50% (208 features; 38,600 reads) and 90% (520 features; 112,600 reads) of samples were stable. One sample did not reach a stable classification during its run, therefore the curves end at 62/63 (98%).

To determine how much sequencing data a stable SteMClass classification requires, we reanalysed each run retrospectively at increasing depth. Of the 66 samples analysed, three were excluded because the final classification score was below 0.5. Stable classification (see Methods) was reached by 62 of 63 included samples. Half of the samples were stable after 208 overlapping CpG features or 38,600 reads, and 90% after 520 features or 112,600 reads (Figure 4c & 4d). The same array- and nanopore-derived methylation data also support CNV profiling to detect culture-acquired aberrations, genotype-based identity verification and sex prediction (Supplementary Figure 9), enabling multiparametric quality control from a single assay. Together, these results provide proof of concept for SteMClass classification and complementary quality control from low pass nanopore sequencing.

### Integrative Broad and CpG-Level Profiling Explains Classification Outcomes

To interpret classification outcomes, we compared validation samples with the reference cohort using the top 20,000 CpGs that distinguish each differentiation state from iPSC (Methods, Supplementary Table 4). Four of the external validation cohort endoderm samples were re-classified as iPSC by SteMClass. Hierarchical clustering of these samples with the reference cohort iPSC and endodermal cells showed clustering with the iPSC samples based on their broader methylation profile (Figure 5a). Top features favouring iPSC over endoderm included the canonical pluripotency marker *POU5F1* and endodermal marker *APC* (Figure 5b). The *POU5F1* promoter showed methylation levels more consistent with reference iPSC in these test samples (Figure 5c). This supports inefficient differentiation rather than misclassification.

**Figure 5.**
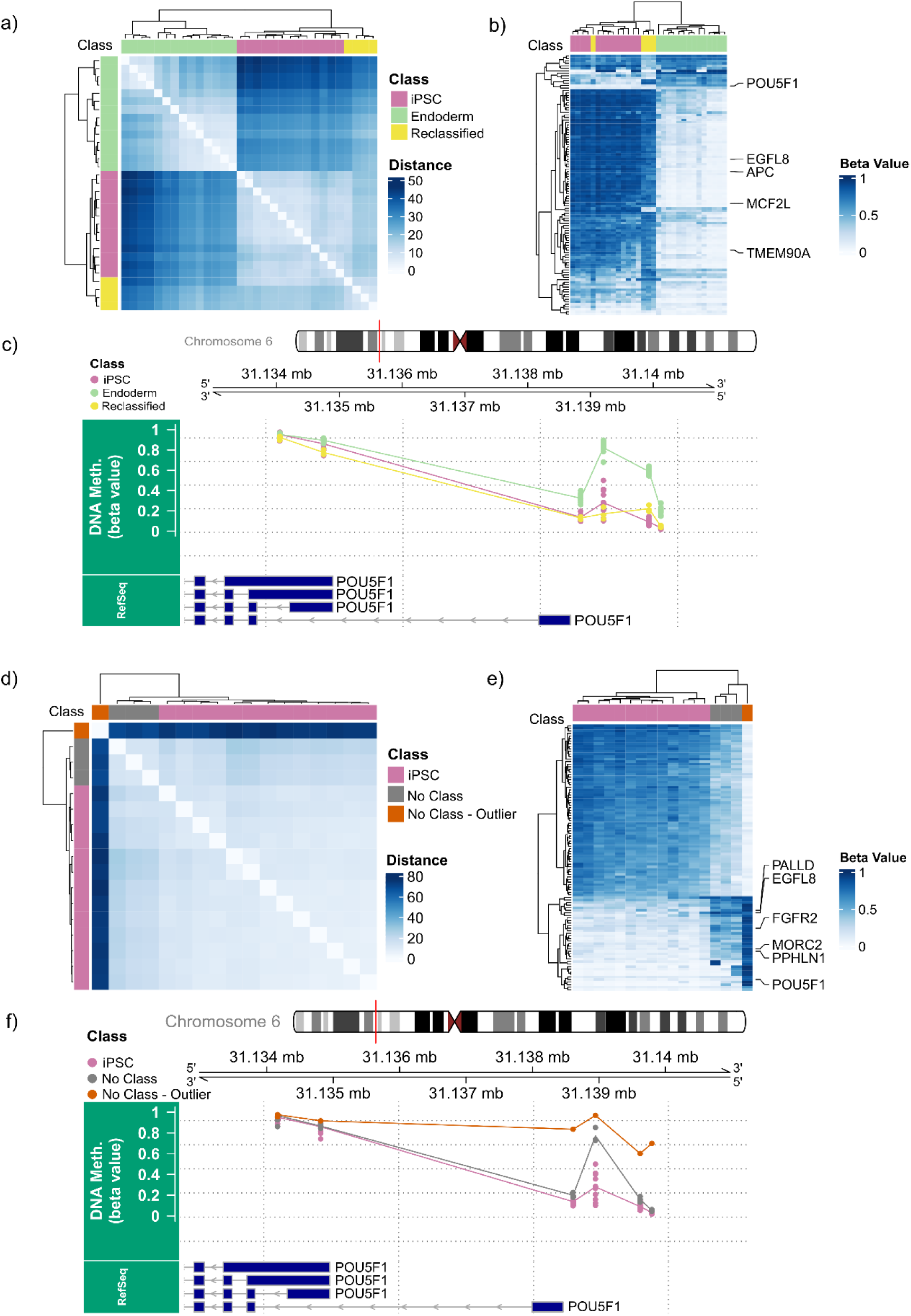
Interpreting SteMClass decisions via broad and narrow view methylation profiling. **(a)** Hierarchical clustering of four samples published as having endoderm differentiation that SteMClass re-classified as iPSC (yellow), together with reference iPSC (pink) and reference endoderm (green) profiles. Clustering was performed on a Euclidean-distance matrix (complete linkage) computed from beta values at the top 20,000 CpGs most discriminating iPSC from all other cell types (“broad view”, Supplementary Table 4). **(b**) Heatmap of the 100 CpGs with highest positive importance for distinguishing iPSC vs. endoderm in the random forest model. Rows represent CpGs and columns represent samples (reference and re-classified endoderm); both were hierarchically clustered using complete linkage with Euclidean distance. Heatmap colours indicate beta value (white = low, blue = high). CpGs associated with canonical markers such as *POU5F1* (pluripotency) and *APC* (endoderm) are labelled. **(c)** Genomic view of beta values across the *POU5F1* promoter region (chr6:31,134,000-31,140,000) for reference iPSC, reference endoderm, and the four re-classified endoderm samples (“narrow view”). Tracks show methylation beta values for each sample (points). **(d)** Hierarchical clustering of four “Not Classifiable” test samples published as iPSC (grey and orange) with SteMClass reference iPSC profiles (pink), clustered as in (a). Of the four rejected samples, three form a distinct clustering branch (grey)with one sample being a clear outlier from all other samples (orange). **(e)** Heatmap of the 100 CpGs whose contributions to the iPSC class were most strongly negative across all four not classifiable samples. **(f)** Genomic view of beta values at the POU5F1 promoter for the four non-classifiable samples.

Four iPSC samples from the external cohort were not classifiable. These four samples formed a cluster distinct from the SteMClass reference iPSCs based on their 20,000 CpG broader methylation profile (Figure 5d). Among the top 100 negatively contributing iPSC features common to these samples were *PALLD* and *FGFR2* which were hypermethylated in promoter CpGs in the four unclassifiable samples (Figure 5e). Three of the samples are more similar to reference iPSCs on the methylation profile, suggesting subtle cumulative differences. One sample exhibited an especially divergent methylation profile, most notably at the *POU5F1* gene promoter, suggesting a questionable undifferentiated state and therefore compromised iPSC quality rather than a classification error for this sample (Figure 5f).

Seven of the eight non-classifiable/reclassified mesoderm samples (all from GSE85828) clustered closer to iPSC and endoderm references than to mesoderm based on the top 20,000 CpG DNA methylation profiles, whereas one sample appeared as a clear outlier across all differentiation states (Supplementary Figure 10a). CpGs driving rejection from mesoderm included mesoderm regulators *MSGN1* and *WNT5B* as well as *APC* and *EFCAB2*. At these sites, the test samples were hypomethylated like the endoderm references relative to the iPSC and mesoderm references (Supplementary Figure 10b). This is in line with the limited epigenetic resolution between mesoderm and endoderm previously reported for this dataset (Schmidt et al., 2023). Among the two non-classifiable ectoderm samples, one was a clear outlier, while the remaining one resembled reference ectoderms in their global profiles (Supplementary Figure 10c & d). Sixteen of twenty external endoderm-annotated samples were not classifiable and all grouped preferentially with iPSC profiles rather than with the highly efficient (>90% CXCR4⁺/SOX17⁺) definitive endoderm reference set (Supplementary Figure 10e). The top 100 CpGs driving rejection from endoderm included lineage-specification loci such as APC (Supplementary Figure 10f). This points to substantial heterogeneity in endoderm differentiation outcomes across protocols rather than misclassification.

#### SteMClass Classification Mirrors Progressive DNA Methylation Shifts During Differentiation

The CpG sites contributing to the non-classifiable predictions for astrocyte samples from the internal (n = 2) and external (n = 2) validation cohorts included three CpGs within the *PAX6* gene body, which ranked among the 100 most informative features for astrocyte classification (Supplementary Figure 10h, Figure 6a). We compared methylation at these CpGs between unsuccessful and successful differentiations (n = 65 samples, Figure 6b).

**Figure 6.**
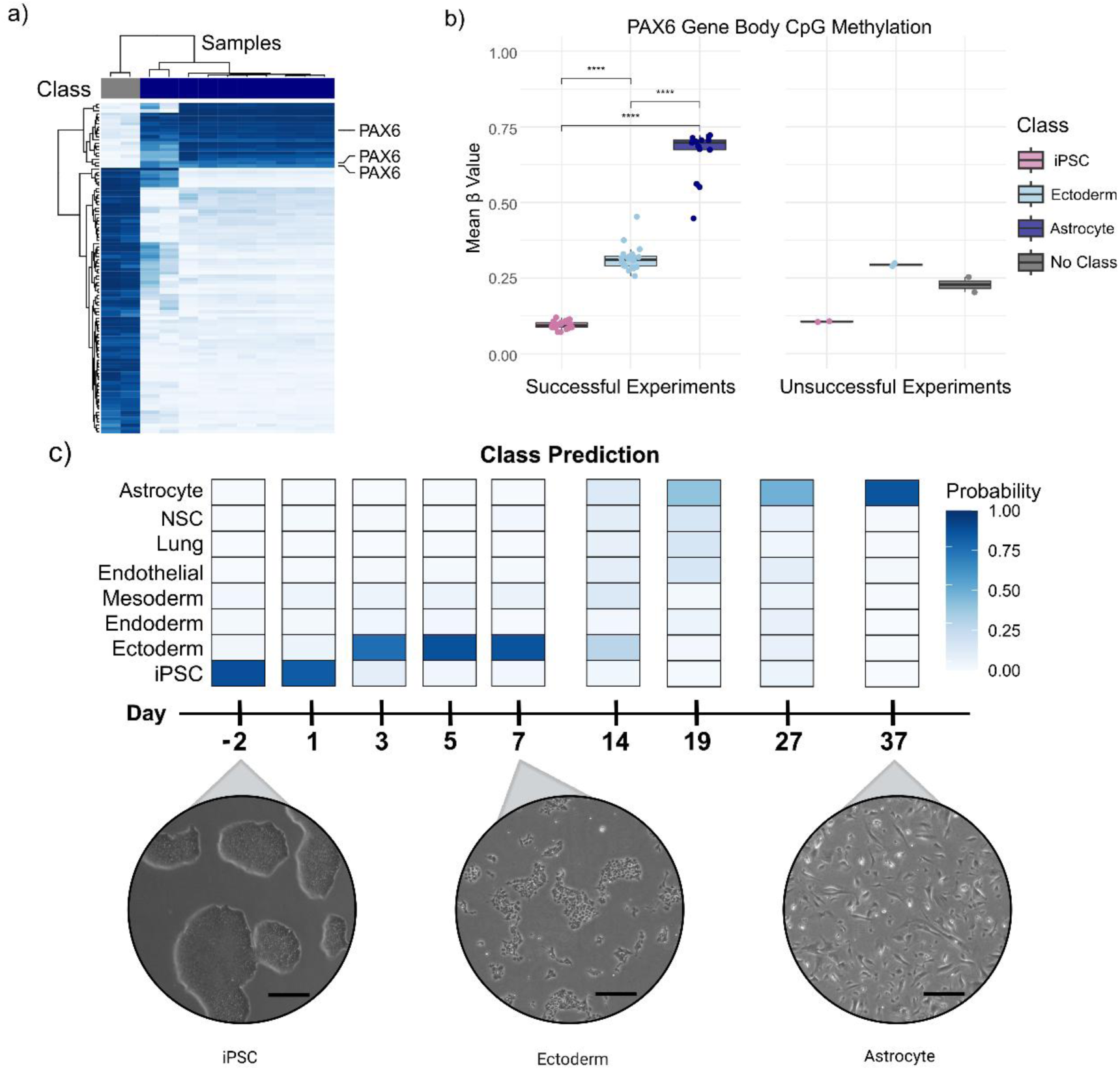
Epigenetic driver inference and dynamic tracking of astrocyte differentiation using SteMClass. **(a)** Heatmap of the 100 CpG sites with the strongest methylation differences (delta beta) in the two not classifiable internal validation samples (grey columns), compared to reference astrocytes (purple columns). Rows represent CpGs and columns represent samples; both were hierarchically clustered using complete linkage with Euclidean distance. Heatmap colour indicates beta value (white = low, blue = high); three *PAX6* gene body-associated CpGs rank among the top features. **(b)** Boxplots show sample-level mean beta-values (0-1) for the *PAX6* CpGs opposing astrocyte classification across iPSC, ectoderm and astrocyte samples. Colours denote SteMClass-assigned differentiation states: iPSC (pink), ectoderm (light blue), astrocyte (dark purple) and “Not classifiable” (grey). Samples are split into panels by differentiation outcome: successful (n = 59) and unsuccessful (n = 6), defined by passing or failing the SteMClass 0.5 cutoff. All pairwise comparisons within the successful group were significant (****, p < 0.0001; Wilcoxon rank-sum). No statistical testing was applied to the unsuccessful group due to limited sample size. **(c)** Temporal SteMClass classification shifts during the same iPSC to astrocyte nine time point differentiation experiment shown in Supplementary Figure 2c. Top panel: heatmap of predicted class probabilities (0-1) at each harvest point for each class represented in SteMClass. Bottom panel: representative culture images at day -2 (classified as iPSC), day 7 (classified as ectoderm), and day 37 (classified as astrocyte), Scale bars = 200 µm.

At the ectoderm stage, methylation in the unsuccessful differentiations was comparable to that of successful experiments (mean β ≈ 0.30), and SteMClass correctly classified these samples as ectoderm. By day 37, however, methylation at these CpGs remained low (mean β ≈ 0.23), rather than increasing towards the levels observed in classifiable astrocytes (mean β ≈ 0.70). Although their scores fell below the 0.5 threshold, the class with the highest score was astrocyte in each case. These findings suggest that unsuccessful differentiations progressed beyond the ectoderm stage but failed to fully acquire the expected astrocyte-associated methylation signature. Classification at intermediate time points could therefore identify deviating differentiations early, allowing them to be discontinued. The three *PAX6* gene-body CpGs provide biologically relevant examples within the top features contributing to astrocyte classification and should not be considered indicators of *PAX6* expression.

Applied to the nine-time-point astrocyte differentiation (Supplementary Figure 2), SteMClass predictions and probability scores shifted progressively as cells moved from pluripotency toward lineage-restricted fates (Figure 6c). The progressive shift in probabilities reflects the model’s sensitivity to intermediate, transient states and its capacity to track subtle changes in lineage commitment.

## Discussion

A fundamental obstacle in stem cell research and regenerative medicine is the lack of standardised frameworks to assess the quality and maturation stage of differentiated cells, causing variability in both results and interpretation. Tools such as the TaqMan hPSC Scorecard and hiPSCore enable assessment of pluripotency and trilineage differentiation potential (Dobner et al., 2024; Lenz, 2012; Tsankov et al., 2015) and more recently DeepDiff added RNA-seq-based classification of developmental states (Dobner et al., 2026). However, these approaches primarily focus on early lineage specification, whereas many downstream applications require differentiation into more specialized cell types that are not comprehensively covered by these assays.

Our prototype classifier, SteMClass, demonstrates strong predictive performance, achieving a cross-validated Brier score of 0.018. Internal and external validation further support its robustness across experimental conditions and its ability to identify potentially problematic samples. Nevertheless, the current reference set does not capture the full diversity of differentiation protocols and classification performance cannot be assumed to generalise to protocols that produce epigenetically distinct differentiation states. Detailed protocol reporting is therefore essential for cross-study comparison. The long-term applicability of SteMClass depends on expanding the reference set with independently validated samples from multiple lines, donors, protocols and laboratories, with retraining and independent validation. We are pursuing this through versioned releases with documented reference compositions and contributions of well-characterised datasets from the community will be essential to extend the classifier across the full range of differentiation protocols and experimental contexts in use.

To preliminarily explore SteMClass performance across different institutions, we incorporated DNA methylation array data for which the published differentiation protocols closely matched our training cohort, allowing for an initial assessment of classifier applicability in varied contexts. The rejection of GSE85828 mesoderm samples aligns with independent analyses of the same dataset, in which the DNA methylation profiles of endoderm- and mesoderm-labelled samples were reported to be almost indistinguishable, in contrast to trilineage-differentiated samples generated in parallel, and in which the magnitude of the endoderm-mesoderm difference was attributed to the differentiation regimen (Schmidt et al., 2023). Consistent with this, mesoderm in GSE85828 was generated using a cardiac differentiation protocol (Zhang et al., 2012) and only the day-5 timepoint was profiled by DNA methylation, whereas the SteMClass mesoderm reference was generated by standardised trilineage differentiation and validated using CD140b (*PDGFRB*, mural/mesenchymal) and CD144 (VE-cadherin, endothelial), markers of lateral plate mesoderm derivatives. These differences in protocol and developmental stage most plausibly explain the outcome, whereas mesoderm samples generated by standardised trilineage differentiation in other studies classified as expected. A similar explanation likely applies to the rejected external endoderm samples. All SteMClass reference endoderm samples were validated by flow cytometry, confirming high differentiation efficiency (>90% SOX17⁺/CXCR4⁺). Previous studies often relied on semiquantitative marker assessments, which have been shown to yield less distinct DNA methylation profiles (Daily et al., 2017; Dobner et al., 2024; Elsafi Mabrouk et al., 2024; Schmidt et al., 2022). For example, GSE253190 reported heterogeneous GATA6 immunofluorescence and SOX17 RT-qPCR without assessing residual pluripotency, while GSE207076 showed persistent OCT4 staining in the representative immunofluorescence images. Collectively, these observations indicate that samples annotated as “endoderm” often represent heterogeneous or incompletely specified states. Rigorously validated, stage-resolved endoderm references could enable discrimination of early regional identities (e.g., anterior versus posterior definitive endoderm) with SteMClass. It has also been shown that iPSC-derived cardiomyocytes frequently show inconsistent gene expression profiles and cell compositions across different laboratories and methods (Grancharova et al., 2021) underscoring the need for more standardised and robust assessment tools.

SteMClass supports evolving DNA methylation profiling technologies, including Illumina Methylation EPIC v2 arrays and nanopore sequencing. Using nanopore-derived methylation profiles, SteMClass correctly classified 63 of 66 samples in our discovery and external-validation cohorts. In an additional cohort spanning pluripotent, trilineage and differentiated cell states, nanopore-based classifications agreed with EPIC-array results for 12 of 13 samples. The same data also support copy number profiling, identity and sex checks, which is particularly relevant for regenerative medicine applications requiring reliable assessment of cell identity and genomic integrity. We routinely multiplex 24 samples across four MinION flow cells, usually reaching our data targets (Figure 4c & d). At approximately half the cost of array profiling, nanopore sequencing offers promising multiparametric quality control, with the potential to identify unsuccessful differentiations at intermediate stages and avoid unnecessary experimental costs. The present nanopore implementation should be regarded as a proof of concept. With coverage of only hundreds of CpGs at shallow depth, a few single-molecule calls can determine whether a sample reaches the threshold. Deeper sequencing or adaptive sampling of classifier CpGs would reduce this variability. Until then, “Not Classifiable” nanopore-based results should prompt deeper sequencing or review rather than be interpreted as definitive. Before nanopore sequencing can fully replace array-based profiling, validation in larger multi-site cohorts and integration of a standardised nanopore workflow into the publicly available SteMClass tool will be required.

While single-cell DNA methylation profiling remains technically complex and expensive, SteMClass’ use of bulk methylation data on homogeneous cell populations provides a practical alternative for cell-type-specific classification.

A promising future direction is extension toward predicting differentiation potential, which would support the screening of iPSC lines for lineage-specific applications relevant to both regenerative medicine and disease modelling. It could also track cell-state composition in sorted organoid and co-culture models. The classifier’s design is a promising base for building QC frameworks GMP compliant to assess stem cell derived Advanced therapy medicinal products (ATMPs). Together, these applications position the classifier as a valuable starting point for advancing stem cell research, quality control in regenerative medicine, and precision medicine applications. In addition to the classification tool itself, the dataset is currently the most extensive DNA methylation data resource of iPSC lines and their isogenic differentiated derivatives across multiple stages of ectoderm, endoderm, and mesoderm, maintained and characterised under standardised conditions and is a resource for studying methylation dynamics during in vitro differentiation.

## Conclusion

SteMClass (https://stemclass.charite.de/) is a publicly available DNA methylation-based classifier prototype that rapidly and reproducibly verifies iPSC and iPSC-derived differentiation states. It identifies and characterises distinct stages of iPSC differentiation, providing a standardised framework for classification that enables consistent cross-laboratory comparisons. As a single, scalable assay with a growing open reference, SteMClass provides a foundation for harmonised quality control of iPSC-derived cells and therefore their clinical translation.

## Supporting information

Supplementary Experimental Procedures

## Acknowledgements

This work was supported by The German Academic Exchange Service (DAAD) [E.P.] and the Berlin School of Integrative Oncology [E.P.]. The work was further supported by the German Cancer Consortium (DKTK), Partner Site Berlin [D.C.]. The authors thank Christine Sers, Gaetano Gargiulio, Frank Heppner and all members of the Heppner lab for helpful discussions and suggestions for this work.

## Author contributions

E.P. performed differentiation experiments, generated DNA methylation data, conducted bioinformatic analyses, prepared figures and tables and wrote the manuscript. V.F.V. performed differentiation experiments, acquisition of nanopore sequencing data and prepared figures and tables. D.T. and P.W. performed DNA extraction and EPIC array processing. C.S. and B.O. contributed to the data processing workflow. P.E. supported nanopore sequencing-based methylation profiling. S.D provided trilineage samples for the nanopore sequencing external validation cohort. D.C., V.F.-V. and H.S. conceived and supervised the study. All authors contributed to manuscript editing and approved the final version.

## Data Availability

Web application source code is available at https://github.com/pereze5/SteMClass_App, all annotation, reference and test data required to reproduce the results reported here are available at https://github.com/pereze5/SteMClass_App/releases and raw data are deposited at GSE308134.

## Declaration of generative AI and AI-assisted technologies in the manuscript preparation process

During the preparation of this work, the author(s) used Claude for improving language and readability. The author(s) reviewed and edited the output as needed and take full responsibility for the content of the published article.

