## Supplementary Figures for "SteMClass: A DNA Methylation-Based Classifier for iPSC In Vitro Differentiation States"

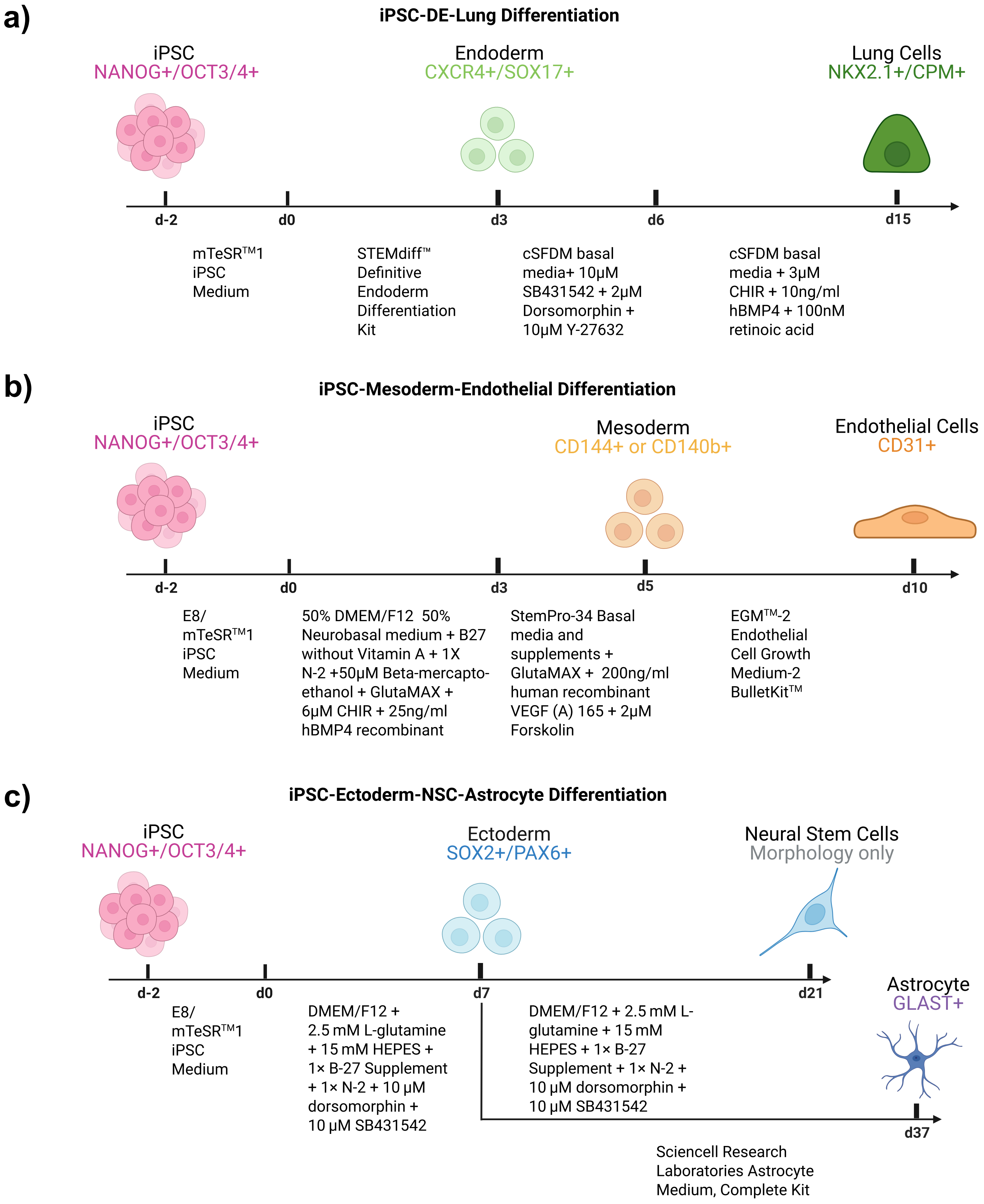


**Supplementary Figure 1.**
Schematic of the iPSC differentiation experiments conducted in this study, including timing of differentiation protocols, growth factor and medium compositions and phentotypic assessment of cell types.

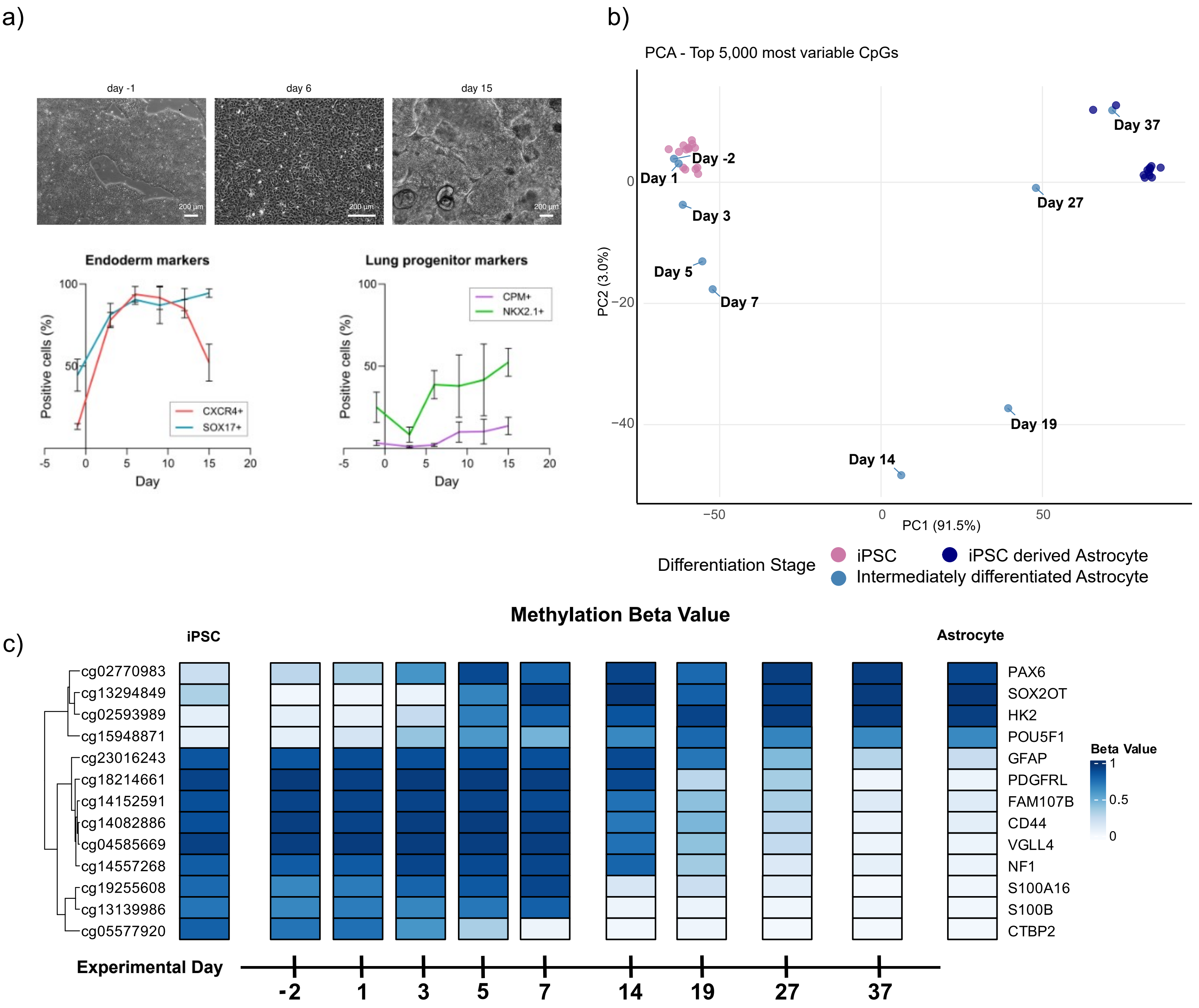


**Supplementary Figure 2.**

**(a)** Top: Representative images during iPSC differentiation to lung. Bottom: Differentiation efficiency assessed by endoderm (CXCR4, SOX17) and lung progenitor markers (Carboxipeptidase-M –CPM- and NKX2.1) using flow cytometry along time. Results are shown as mean +/- SD (n=4 biological replicates; BIHi001-B, BIHi005-A, BIHi250-A and UCSFi001-A). **(b)** Principal component analysis (PCA) plot of DNA methylation beta values from the 5,000 most variable CpG sites (selected by across-sample standard deviation) in iPSCs (n = 14) and iPSC-derived astrocytes (n = 12) (two parental iPSC lines), together with nine intermediate time-point samples from the differentiation time course experiment. Data were mean-centred and scaled prior to PCA. Each point denotes an independent technical replicate (separately cultured and processed). Colours indicate differentiation states. PC1 (91.5% variance) separates iPSCs from astrocyte derivatives and intermediate states, reflecting progression along the differentiation axis. **(c)** DNA methylation dynamics were evaluated during an individual 37-day iPSC derived astrocyte differentiation experiment. For visualisation purposes, the 13 CpG sites showing the largest differences in beta values between undifferentiated and differentiated states are depicted. The heatmap displays mean beta values across iPSC samples (n = 14), individual beta values at the nine intermediate differentiation experiment stages, and mean beta values in astrocyte samples (n = 12). Rows correspond to CpG sites, ordered by hierarchical clustering using Pearson correlation distance. CpG loci are annotated with CpG unique IDs and their associated gene names.

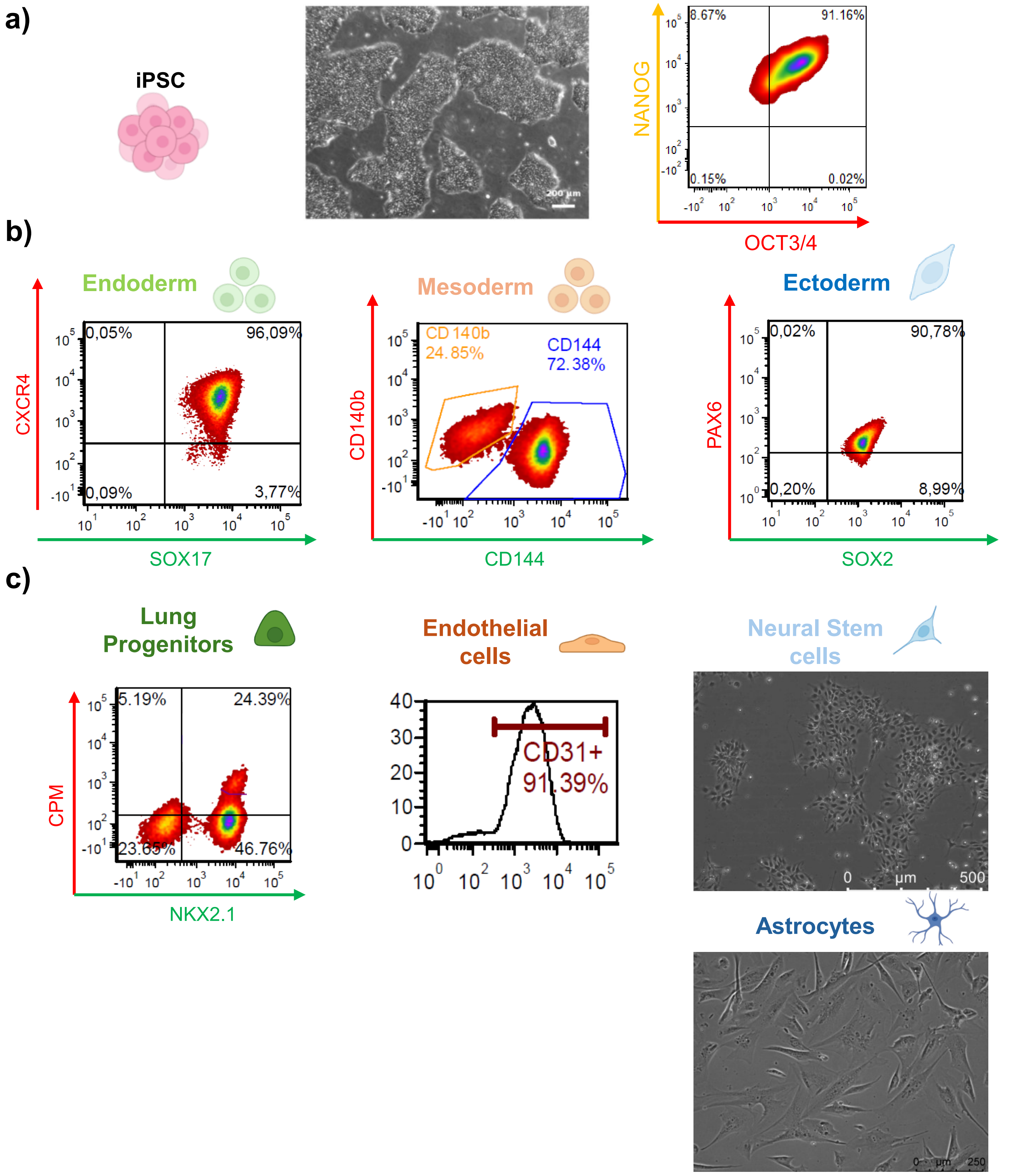


**Supplementary Figure 3.**

Representative quality control performed on samples confirmed iPSC and iPSC-derived differentiated cells phenotype and purity. **(a)** Representative image of iPSC morphology and dotplot showing co-expression of undifferentiated markers determined by FACS in BIHi250-A line. Scale bar: 200µm. **(b)** Representative dotplots showing the expression of specific markers used to evaluate differentiation of iPSC into three germ layers determinedby FACS. Example belongs to trilineage differentiation performed on BIHi250-A line. **(c)** Examples of specific marker expression and/or morphology of iPSC derived differentiated cells from: Lung progenitors d15 (BIHi001-B, scale bar: 200µm), endothelial cells d11 (BIHi005-A, scale bar: 100µm), neural stem cells d21 (BIHi001-B,scale bar: 500µm) and astrocytes d30 (BIHi001-B, scale bar:250µm). Heterogeneous populations, here depicted by lung progenitors, were enriched by FACS using specific marker(e.g. CPM+) prior methylation status evaluation. Astrocytes were MACS purified for GLAST positive cells, NSCs were characterised at day 7 by FACS and day 18-21 by morphology..

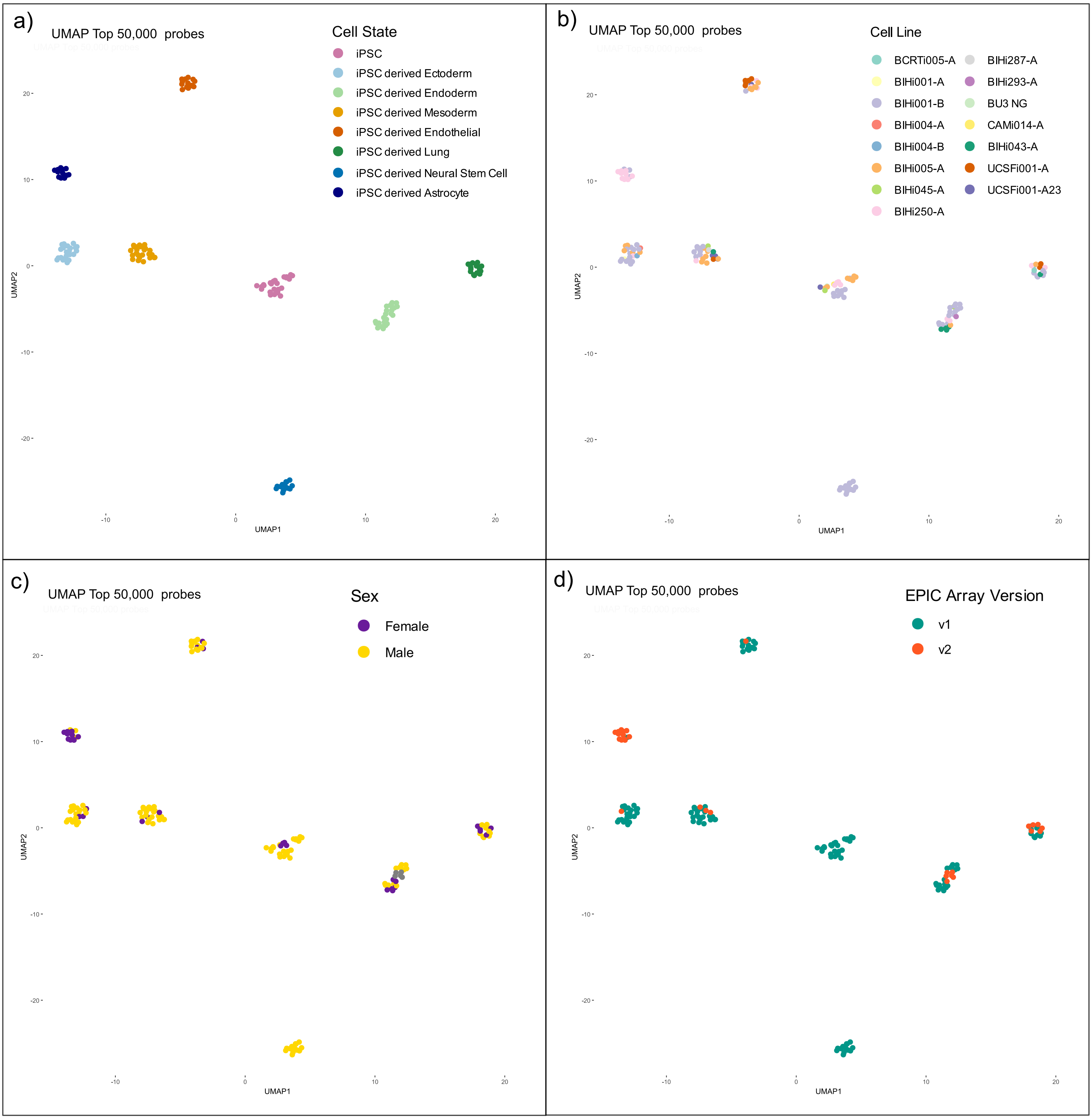


**Supplementary Figure 4 a-d.** UMAP projections of the full dataset illustrating that the separation of the eight cell-state classes is not impacted by common confounders. **(a)** Points coloured by reported cell phenotype ("cell type") show distinct clusters. **(b-d)** The same embedding is coloured by potential batch or biological covariates: **(b)** donor cell line, **(c)** donor sex, and **(d)** EPIC array version. The preservation of the cell-type structure across panels despite variation in these factors indicates that the observed clustering is independent of these potential batch effects.

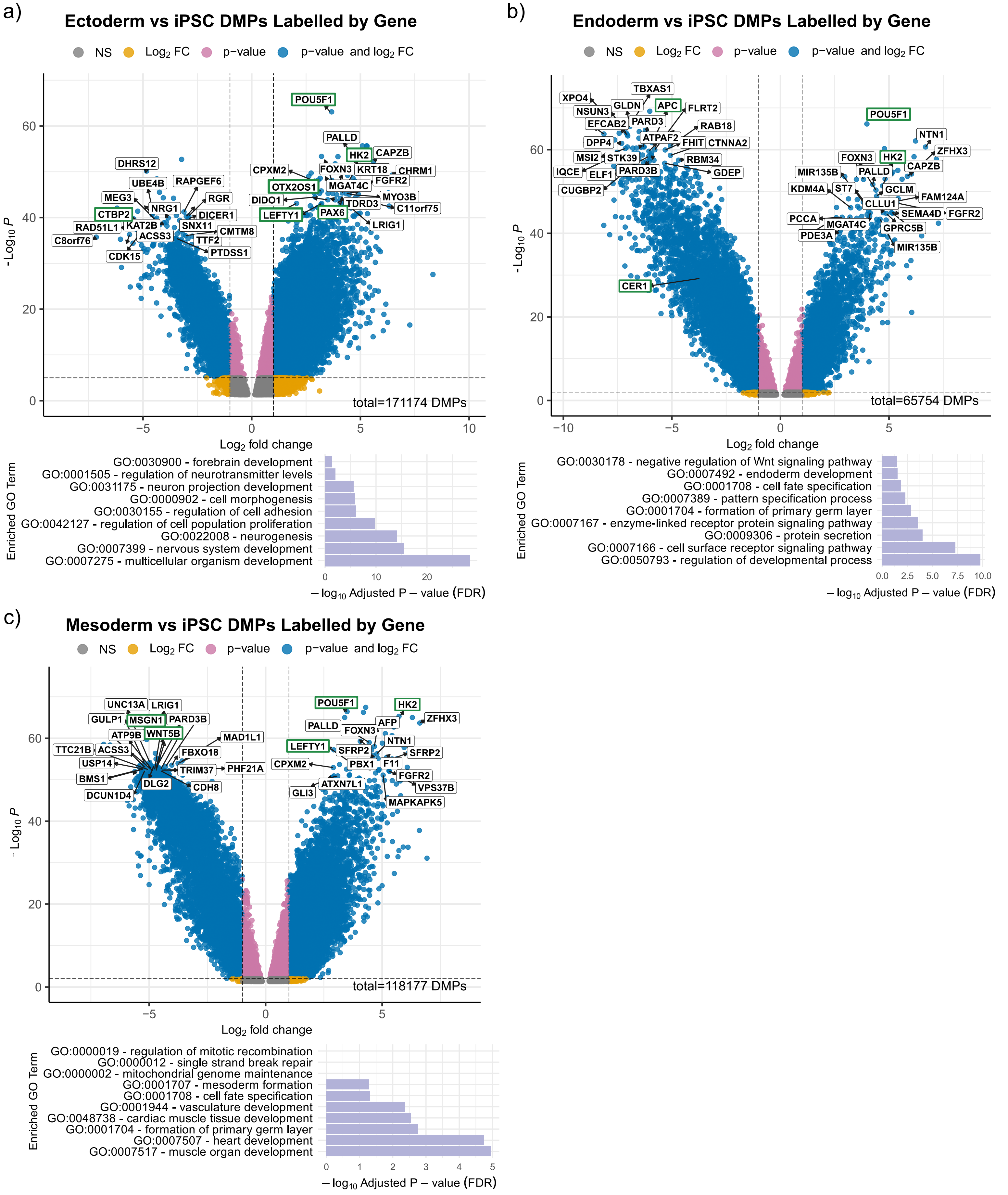


**Supplementary Figure 5.**
**(a-c)** Volcano plots of differentially methylated positions (DMPs) comparing iPSCs to: **(a)** ectoderm, **(b)** endoderm, **(c)** mesoderm. Each point is a differentially methylated CpG; blue points meet significance thresholds (FDR-adjusted $p < 0.05$ and $|\log_2 \text{fold change}| > 1.5$). The most significant CpGs are annotated with their associated gene. Canonical or candidate novel lineage markers are highlighted in green to emphasize key methylation shifts underlying differentiation. Bar plots summarize gene ontology enrichment among significant DMP-associated genes, showing overrepresented biological processes and pathways consistent with the expected differentiation programs. NS, not significant.

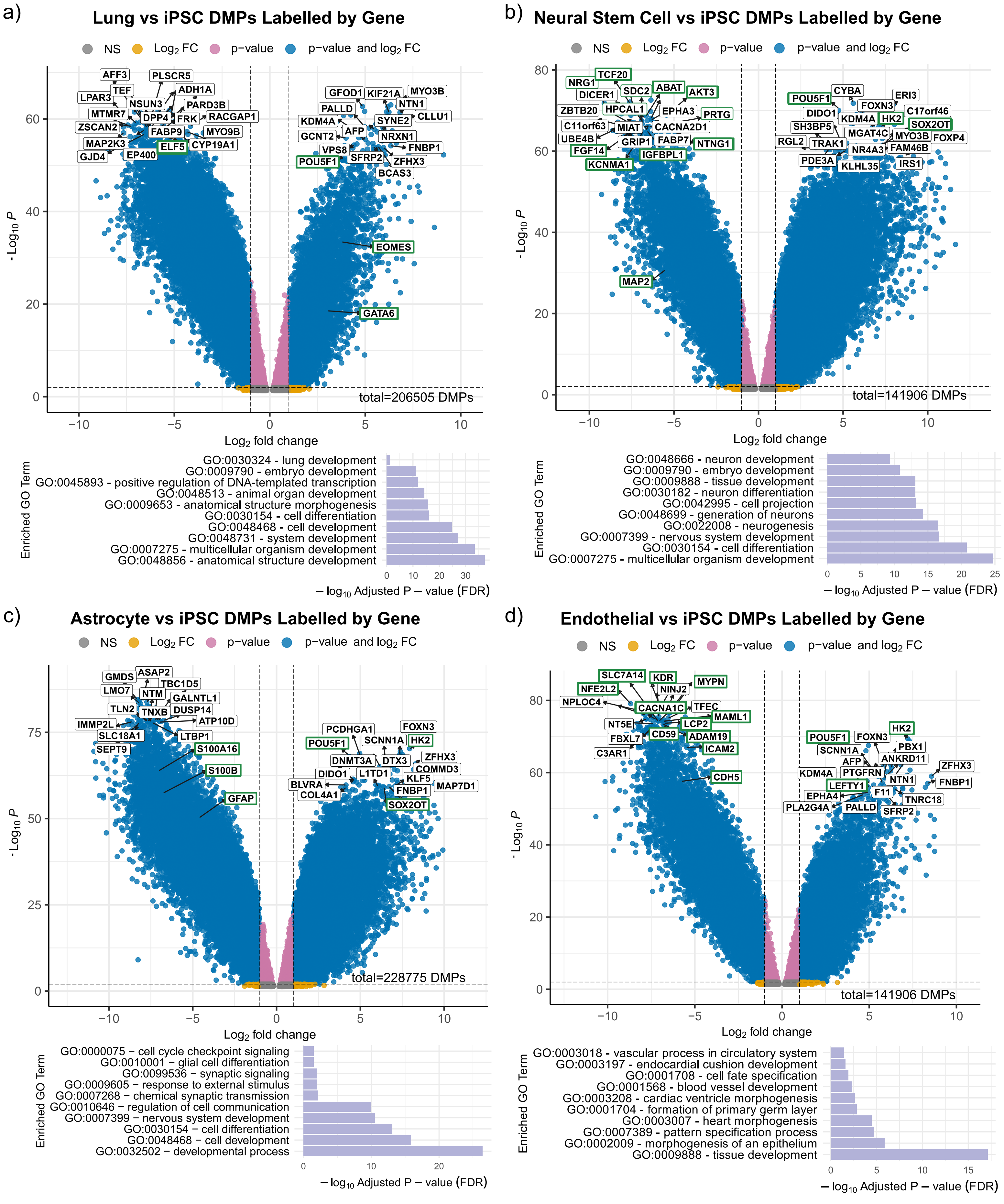


**Supplementary Figure 6.**
**(a-d)** Volcano plots of differentially methylated positions (DMPs) comparing iPSCs to: **(a)** lung cells, **(b)** neural stem cells, **(c)** astrocytes, **(d)** endothelial cells. Each point is a differentially methylated CpG; blue points meet significance thresholds (FDR-adjusted $p < 0.05$ and $|\log_2 \text{fold change}| > 1.5$). The most significant CpGs are annotated with their associated gene. Canonical or candidate novel lineage markers are highlighted in green to emphasize key methylation shifts underlying differentiation. Bar plots summarize gene ontology enrichment among significant DMP-associated genes, showing overrepresented biological processes and pathways consistent with the expected differentiation programs. NS, not significant.

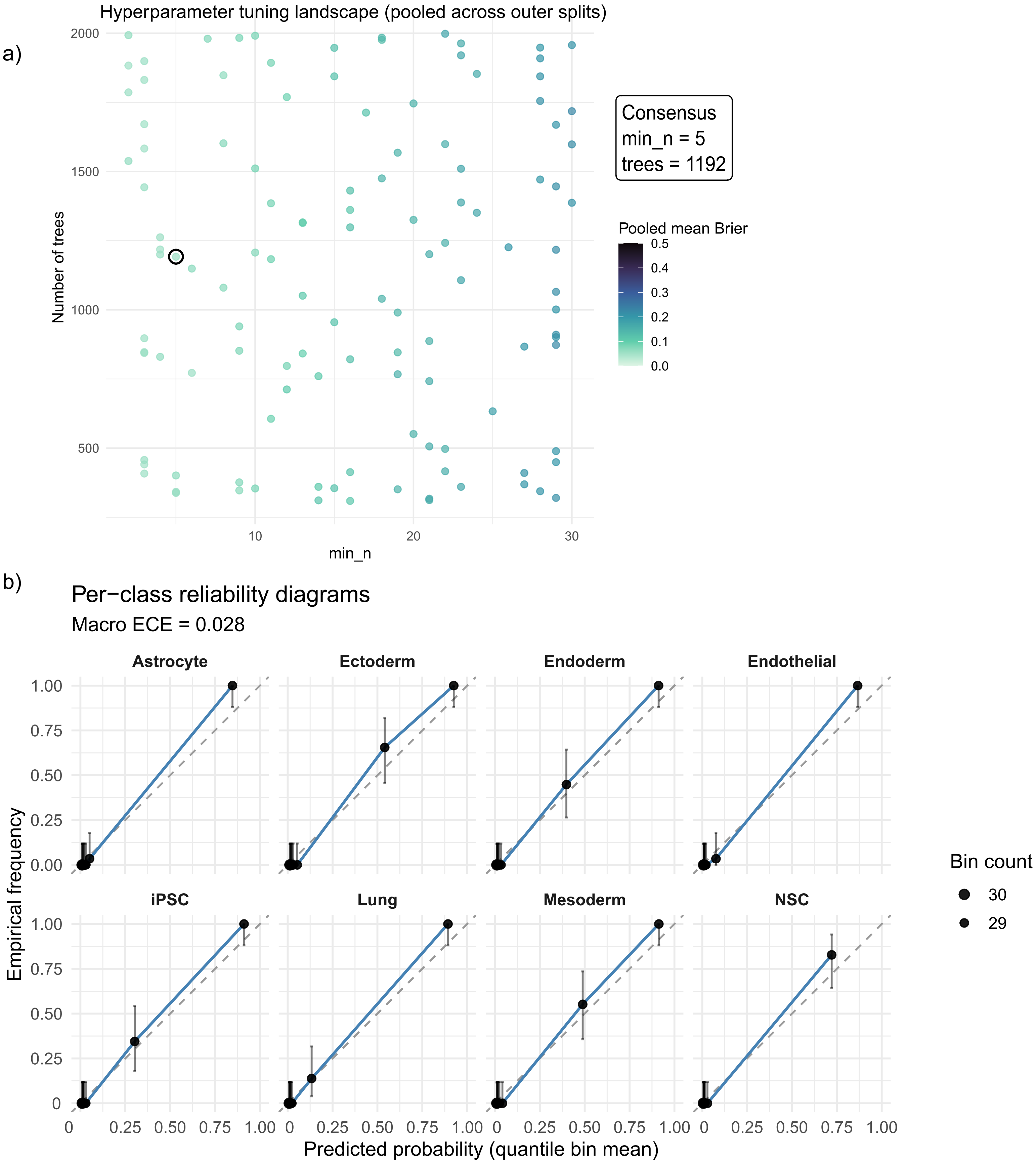


**Supplementary Figure 7.**
**(a)** Hyperparameter tuning landscape pooled across outer resamples. Points show candidate min_n and tree combinations evaluated by inner cross-validation, coloured by the outer-split averaged multiclass Brier score. Black circle denotes the consensus setting used for final training. **(b)** Per-class reliability diagrams for the eight cell-state classes based on pooled out-of-fold calibrated predictions. For each class, predicted probabilities were grouped into deciles using equal-frequency (quantile) binning, and the mean predicted probability within each bin was plotted against the empirical class frequency (one-vs-rest). The dashed diagonal denotes perfect calibration. Vertical error bars show exact binomial 95% confidence intervals, and point size reflects the number of samples per bin. The subtitle reports the macro expected calibration error (ECE), defined as the mean across classes of the bin count-weighted absolute difference between predicted probability and empirical frequency. These plots assess probabilistic calibration across the prediction range.

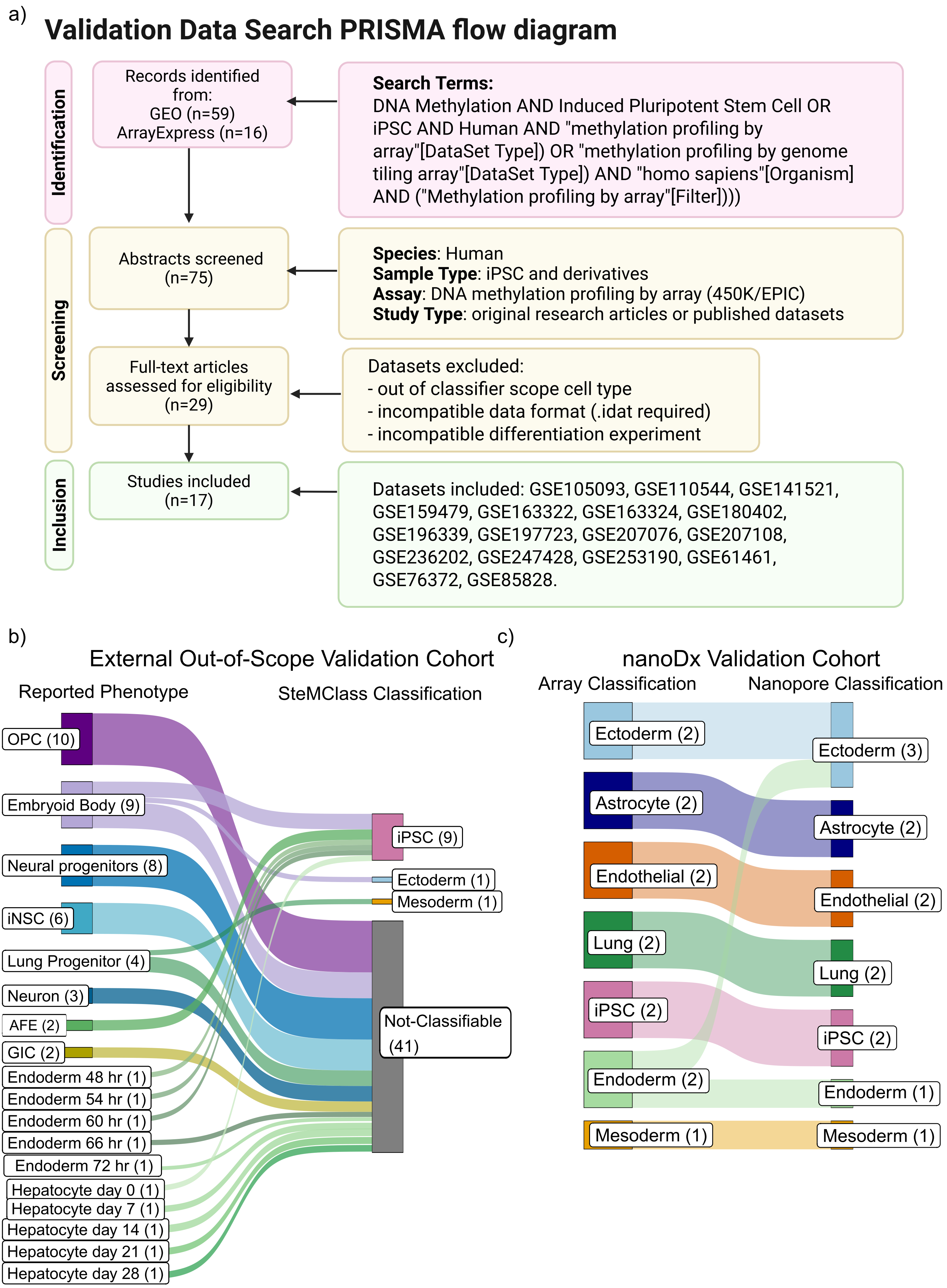


**Supplementary Figure 8. Multi-Cohort and Cross-Platform Validation of SteMClass.**

**Supplementary Figure 8. Multi-Cohort and Cross-Platform Validation of SteMClass.**
**(a)** External validation cohort assembly. PRISMA-style flow diagram summarizing the systematic curation of publicly available DNA methylation array datasets: records identified, screened, included, and excluded with reasons. A total of 249 samples were assembled from independent laboratories; based on the original publications' methods and annotations, these samples were considered possibly comparable to the eight cell-state classes defined in this study. Raw signal intensity files (.IDAT) were downloaded from GEO and processed uniformly. This external cohort was intended to evaluate SteMClass performance on independently generated data and to evaluate the variability of characterisation in the literature. **(b)** Sankey diagram comparing reported phenotype to SteMClass predictions for the external Out-of-Scope cohort (n = 54) assembled from publicly available methylation datasets. Flows connect the original sample annotation to the assigned class (including "Not Classifiable" rejections). Samples were those whose metadata indicated cell states not currently represented by SteMClass. Six early-stage (< 3 day) endoderm samples and three embryoid body samples were classified as iPSC, one embryoid body sample as ectoderm, and one lung progenitor (day 25) sample as mesoderm, while all remaining samples were rejected. **(c)** Sankey diagram comparing SteMClass classifications on EPIC array data to classifications on Nanopore methylation calls for a subset of the internal hold-out test set (n = 13). Each flow connects the class assigned by SteMClass on array-derived beta values (left) to the class assigned by an ad-hoc Random Forest model applied to Nanopore methylation calls via the nanoDx pipeline (right). Twelve of thirteen samples are classified identically on both platforms, with a single discordant call highlighted.

a)

**Nanopore Sequencing**

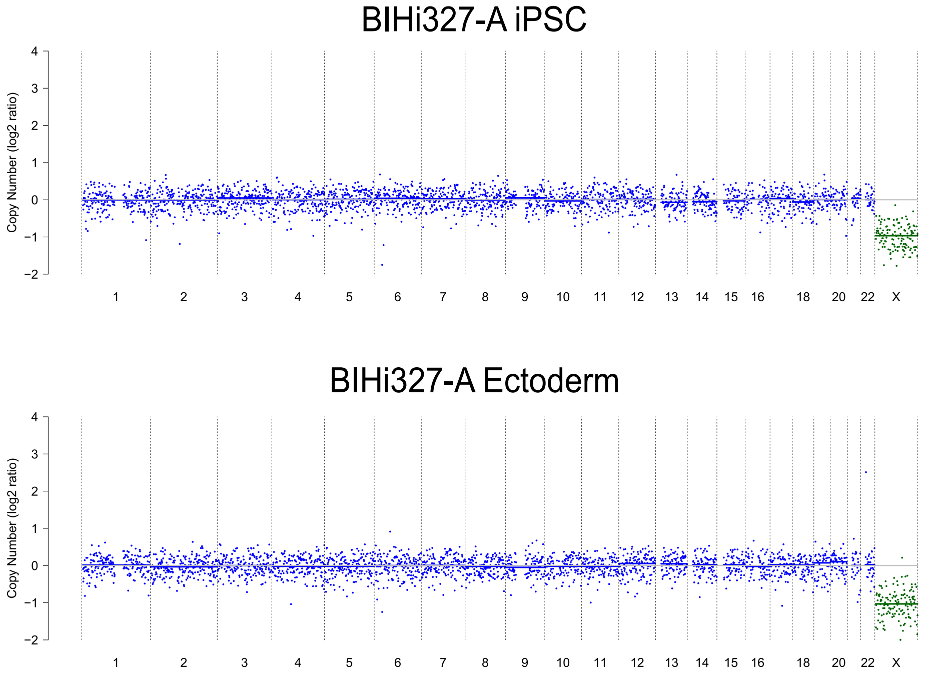


b)

**Infinium MethylationEPIC v2.0 Array**

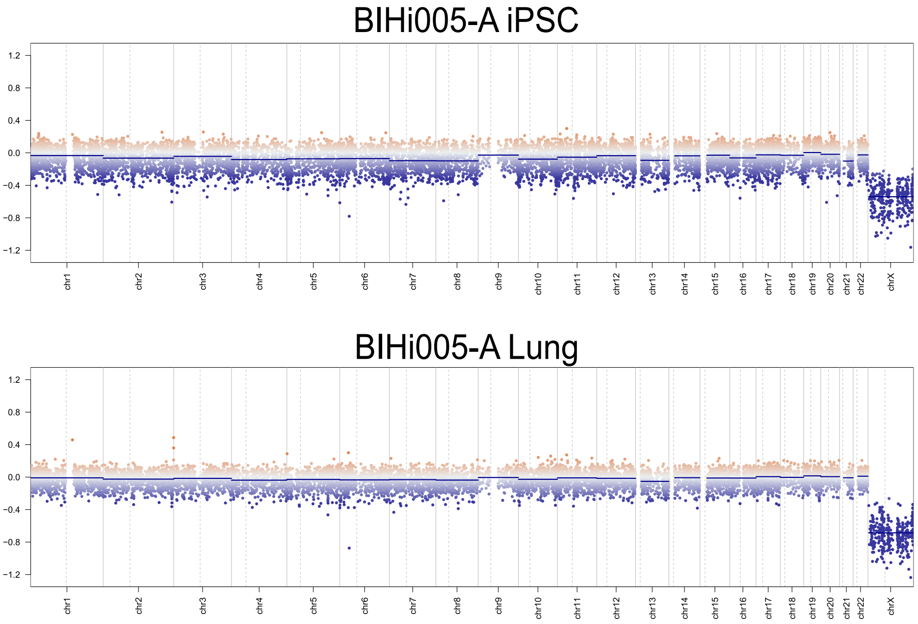


**Supplementary Figure 9. Copy number profiles of iPSCs before and after differentiation inferred from nanopore sequencing and methylation array data.**

**(a)** Genome-wide copy number profiles from nanopore sequencing of an iPSC sample before differentiation (top) and after 7 days of ectoderm differentiation (bottom). Copy number profiling was performed with ichorCNA v0.5.1 using 1 Mb bins. **(b)** Genome-wide copy number profiles from Infinium MethylationEPIC v2.0 array data of an iPSC sample before differentiation (top) and after 15 days of lung differentiation (bottom). Copy number analysis was performed with the conumee 2.0 Bioconductor package v2.1.2. In all panels, chromosomes are shown along the x-axis and log2 copy number ratios on the y-axis. Points represent bins (nanopore) or probes (EPICv2), vertical lines mark chromosomal boundaries and dashed lines mark the centromeres. No copy number alterations were acquired during differentiation. Because a female reference set was used, male samples show an apparent loss of chromosome X, which reflects sex rather than a copy number alteration.

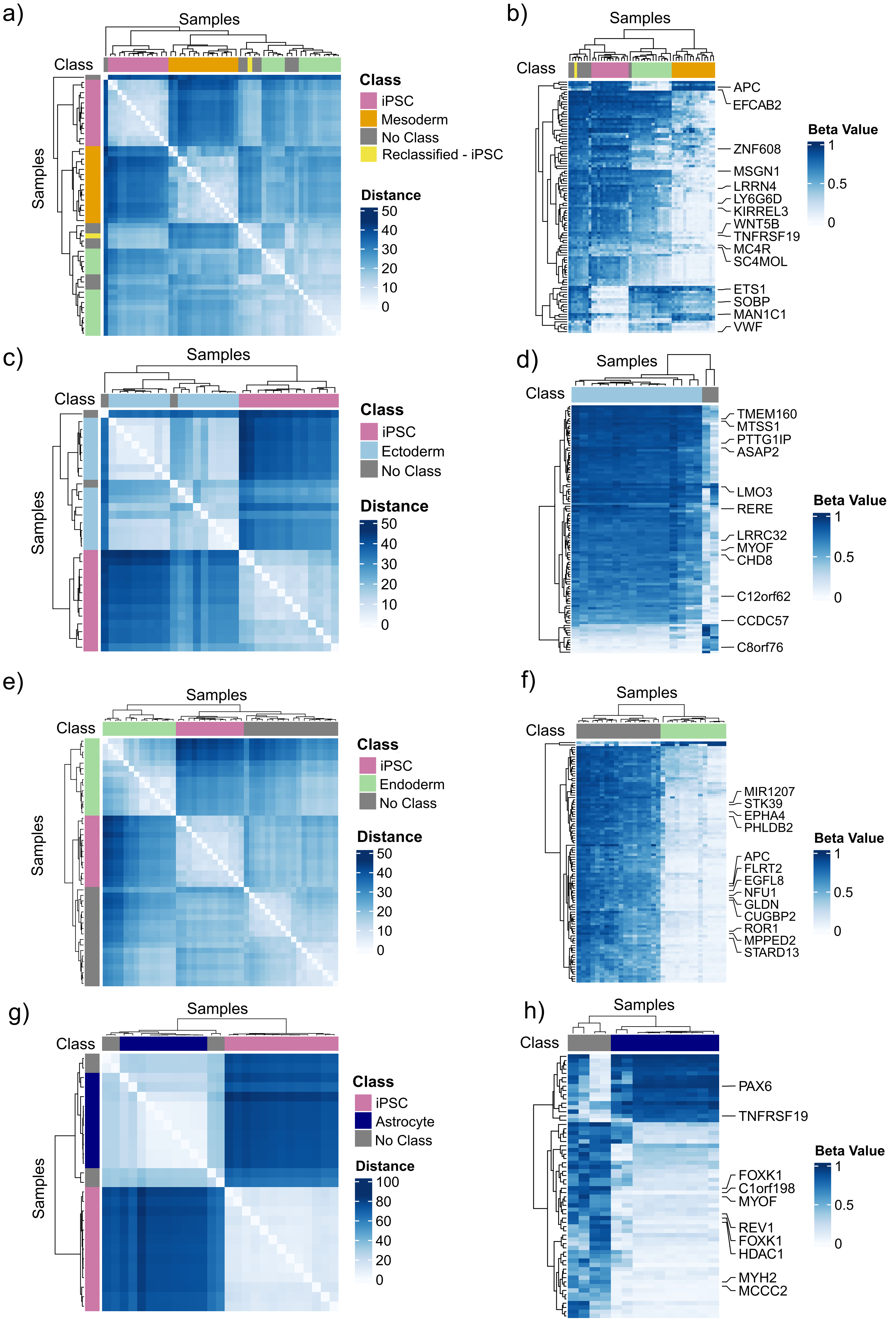


**Supplementary Figure 10. Interpreting SteMClass decisions via broad and narrow view methylation profiling (continued).**

**Supplementary Figure 10. Interpreting SteMClass decisions via broad and narrow view methylation profiling (continued).**
**(a)** Hierarchical clustering of seven mesoderm-annotated samples that SteMClass rejected (grey), one that was reclassified as iPSC (yellow) alongside reference mesoderm (orange), reference endoderm (green) and reference iPSC (pink) profiles, using beta values at the top 20,000 mesoderm lineage-distinguishing CpGs (Supplementary Table 4). **(b)** Heatmap of the 100 CpGs whose contributions to the mesoderm class were most strongly negative across the eight non-classifiable/reclassified mesoderm samples. Rows are CpGs, columns are samples, and heatmap colour indicates beta value (white = low, blue = high). CpGs in canonical mesoderm regulators such as MSGN1 and WNT5B are labelled. **(c)** Hierarchical clustering of two ectoderm-annotated samples that were rejected by SteMClass (grey) with reference ectoderm (blue) and iPSC (pink) profiles, using the iPSC specific 20,000-CpG panel. One sample (GSM2285110) is a clear methylation outlier, while the other one clusters near reference ectoderms. **(d)** Heatmap of the 100 CpGs with strongest negative contribution to ectoderm classification across the seven rejected samples. **(e)** Hierarchical clustering of eighteen endoderm-annotated samples rejected by SteMClass (grey) against reference endoderm (green) and iPSC (pink) profiles. All rejected endoderm samples cluster closer to iPSCs. **(f)** Heatmap of the 100 CpGs whose contributions most strongly oppose endoderm classification in the rejected samples. APC-associated CpGs rank among the top features. **(g)** Hierarchical clustering of four astrocyte annotated samples rejected by SteMClass (grey) against reference astrocyte (purple) and iPSC (pink) profiles. All rejected astrocytes display distinct broad-view DNA methylation profiles when compared to the reference cohort. **(h)** Heatmap of the 100 CpGs whose contributions most strongly oppose astrocyte classification in the rejected samples. Three PAX6-associated CpGs rank among the top features.
