## Supplementary Experimental Procedures for "SteMClass: A DNA Methylation-Based Classifier for iPSC In Vitro Differentiation States"

### Supplemental Experimental Procedures

#### Cell Culture and Differentiation Experiments

##### Culture of iPSCs

Human induced pluripotent stem cells (iPSCs) were maintained on growth factor reduced Geltrex (Gibco) in E8 or mTeSR (Stem Cell Technologies) media with daily media exchange. Cells were enzymatically clump-passaged when >70% confluency was reached, usually every 2-4 days using 0.5mM EDTA (UltraPure™ EDTA, Thermo Fisher). Supplementary Table 1 compiles the iPSC lines and subclones used in this study and their maintenance culture conditions, detailed protocol information is available on Protocols.io (Fernandez Vallone et al., 2025). All iPSC lines and subclones were provided by Berlin Institute of Health Core Unit Pluripotent Stem Cells and Organoids (BIH-CUSCO) and have ethical approval for their use. All iPSC lines and subclones used in this study were tested for genomic stability as reported in Supplementary Table 1. Briefly, for G-banding mitotic arrest was induced with Colcemid and metaphases were analysed at a minimum resolution of 200 bands per haploid chromosome set. For all cases 20 metaphases were counted. Copy number variations (CNVs) were assessed via SNP arrays (OmniExpressExome BeadChip, Illumina) on DNA samples (min 200ng). Analysis was performed using an in-house developed pipeline (Kügelgen et al., 2025). Acquisition of de novo CNVs was flagged by comparison with reference sample. Results for both genomic stability tests are available upon request.

##### Differentiation Protocols

Cell type nomenclatures, iPSC lines and corresponding differentiation protocols used in this study are described in Supplementary Tables 1-3 and Supplementary Figure 1.

##### Differentiation to Ectoderm

iPSCs were dissociated into single cells using TrypLE (Gibco) and seeded at a density of $5 \times 10^5$ cells per well in a 6-well plate ($5.2 \times 10^4$ cells/cm$^2$) coated with 1:120 dilution of Geltrex™ (Gibco) in Essential 8™ Medium supplemented with 10 µM Y-27632 (Selleck Chemicals). Cells were cultured for 2 days with daily medium changes using Essential 8™ Medium. After 2 days, the medium was replaced with neural induction medium (NIM) promoting dual SMAD inhibition. NIM was composed of DMEM/F12 supplemented with 2.5 mM L-glutamine, 15 mM HEPES, 1× B-27 Supplement, 1× N-2 Supplement (all from Thermo Fisher Scientific), 10 µM dorsomorphin (Tocris Bioscience), and 10 µM SB431542 (Tocris Bioscience) (Supplementary Figure 1c). Medium was replenished every 24 hours for 7 days, after which cells were characterised using fluorescence-activated cell sorting (FACS) to assess the expression of the markers PAX6 (Miltenyi Biotec, 130-123-328), Nestin (Miltenyi Biotec, 130-119-799), SOX1 (Miltenyi Biotec, 130-111-042), and SOX2 (Miltenyi Biotec, 130-120-721). Samples with >85% positive expression for these markers were harvested and stored as cell pellets at -80 °C. Ectoderm day 7 stage was used as starting point to differentiate Neural Stem Cells and Astrocytes.

**Differentiation to Neural Stem Cells**

iPSCs were differentiated into ectodermal cells as described above for 7 days, after which cells were analysed by FACS for the markers PAX6 and SOX2 (Supplementary Figure 1c). Batches with >80% marker expression were dissociated into single cells and seeded at a density of approximately 15,000 cells/cm$^2$ onto plates coated with either Geltrex™/Matrigel® (1:120 dilution in DMEM), or vitronectin (1:100 dilution in PBS without $Ca^{2+}$, $Mg^{2+}$) in Neural Induction Medium composed of DMEM/F12 supplemented with 2.5 mM L-glutamine, 15 mM HEPES, 1× B-27 Supplement, 1× N-2 Supplement (all from Thermo Fisher Scientific), 10 µM dorsomorphin (Tocris Bioscience), and 10 µM SB431542 (Tocris Bioscience). Medium was replenished every 24 hours for 18 days, after which cells were assessed morphologically, harvested and stored as pellets at -80 °C.

**Differentiation to Astrocytes**

Astrocyte progenitor cells were generated from iPSCs as previously described (TCW et al., 2017) (Supplementary Figure 1c). Briefly, iPSCs were differentiated into ectodermal cells as described above for 7 days, then dissociated into single cells and seeded at a density of 15,000 cells/cm$^2$ onto plates coated with Geltrex™/Matrigel® (1:120 dilution in DMEM), or vitronectin (1:100 dilution in PBS without $Ca^{2+}$, $Mg^{2+}$) in Neural induction medium supplemented with 10 µM Y-27632. Medium was changed every 24 hours for 48 hours, followed by replacement with complete Astrocyte Medium (ScienCell Research Laboratories, Cat# 1801). Medium was changed every 48 hours, and when cells reached 90-95% confluency (every 6-7 days), they were passaged to the initial seeding density of 15,000 cells/cm$^2$. After 30 days, cells were dissociated, and GLAST-positive cells were isolated using the Anti-GLAST (ACSA-1) MicroBead Kit and a MiniMACS™ Separator (Miltenyi Biotec, 130-095-825), following manufacturer´s instructions. Isolated GLAST-positive cells were stored as pellets at -80 °C.

**Differentiation to Endoderm**

Before endoderm differentiation iPSC lines which were not cultured in mTeSR™1 (Stem Cell Technologies) on Geltrex (Gibco) coated plates, were adapted for at least 2 passages to these conditions. 70-80% confluent iPSC were harvested as single cells using Accutase (Gibco). Three million cells per well of 6WP were seeded on Geltrex coated surface in mTeSR™1 supplemented with 10µM Y-27632 (Stem Cell Technologies). After 24 hrs, endoderm differentiation was started using the STEMdiff™ Definitive Endoderm Differentiation Kit (Stem Cell Technologies) following manufacturer´s instructions. This day was considered day 0. At day 3 cells were harvested using Accutase and immunostained for FACS analysis using CD184

(CXCR4) (Miltenyi Biotec, 130-120-708) and SOX17 (Miltenyi Biotec, 130-111-031). Definitive endoderm differentiation with >90% efficiency were used to derive lung progenitors and a subfraction of the cells were stored at -80°C as a pellet for DNA isolation and downstream application.

**Differentiation to Lung Cells**

iPSC derived definitive endodermal cells were derived as described above and further differentiated to lung progenitors according to Jacob et al. (Jacob et al., 2019) (Supplementary Figure 1a). Briefly, day 3 endoderm cells harvested as clumps using Gentle Dissociation reagent (Stem Cell Technologies) were seeded in 6WP as 1:4 and 1:6 ratios in cSFDM basal medium (described in Jacob et al., 2019) supplemented with 10μM SB431542 (Sigma), 2μM Dorsomorphin and 10μM Y-27632 (Stem Cell Technologies). After 24 hrs, medium was changed removing Y-27632 (day 4). Anterior foregut differentiation continued another 48hs without media change. At day 6, medium was changed to lung progenitor induction medium conformed by cSFDM basal medium supplemented with 3μM CHIR (EZSolution Avantor), 10ng/ml hBMP4 (Prepotech) and freshly added 100nM retinoic acid (Sigma). Medium was changed every 48 hrs until day 15. Lung progenitors were isolated based on Carboxypeptidase M (CPM) expression using anti human-CPM antibody (WAKO Fujifilm, 014-27501) and subsequent magnetic separation using Anti-mouse IgG2a+b Microbeads (Miltenyi Biotec, 130-047-201), MACS separation columns (Miltenyi Biotec, 130-042-201) and MiniMACS Separator (Miltenyi Biotec). CPM positive cells were stored as cell pellet at -80°C.

**Differentiation to Mesoderm: Endothelial and Pericyte Progenitor Cells**

Mesoderm and subsequent endothelial cell differentiation was adapted from Patsch et al. (Patsch et al., 2015) (Supplementary Figure 1b). Briefly, 70-80% confluent iPSC were harvested as single cells using Accutase (Gibco). Four hundred thousand cells were seeded per well of 6WP on Geltrex coated surface in E8 or mTeSR media (hiPSC line dependent) supplemented with 10μM Y-27632 (Stem Cell Technologies). After 24 hrs, at day 0, medium was exchanged for 4ml per well of Endo Priming medium composed by 50% DMEM/F12 (Thermo Fisher, 11320-033) and 50% Neurobasal medium (Thermo Fisher, 21103049) supplemented with B27 without Vitamin A (Gibco), N-2 (Gibco), 50μM Beta-mercapto-ethanol (Gibco), GlutaMAX (Gibco), 6μM CHIR (EZSolution Avantor), and 25ng/ml hBMP4 recombinant (Prepotech). No medium change was performed for 72 hrs. At day 3, medium was exchanged for Endo Induction medium consisting of StemPro-34 Basal medium and supplements (Gibco), GlutaMAX (Gibco), 200ng/ml human recombinant VEGF (A) 165 (Prepotech) and 2μM Forskolin (Abcam). Cells were cultured for 48 hrs. At day 5, cells were harvested as single cells using Accutase (Gibco). CD144 was used as surface marker to assess differentiation efficiency (Miltenyi Biotec, 130-100-742) by FACS and to isolate endothelial progenitors using magnetic MicroBeads (Miltenyi Biotec, 130-097-857), MACS separation columns (Miltenyi Biotec, 130-042-201) and MiniMACS Separator (Miltenyi Biotec), following manufacturer´s instructions. Cells belonging to the CD144 positive and negative fraction (Mesoderm) were kept as pellets at -80°C. Cells belonging to the CD144 positive fraction were used as the starting population for endothelial cell differentiation. Cells of the negative fraction are considered pericyte progenitor cells.

##### Differentiation to Endothelial Cells

Endothelial Progenitor cells (CD144+) obtained as described above were seeded on Gelatin 1% (Millipore) coated T25 flasks in complete EGM$^{TM}$-2 Endothelial Cell Growth Medium-2 BulletKit$^{TM}$ (Lonza). Between day 12-14 when cells were 90% confluent, they were harvested as single cells using Acutasse (Gibco). FACS for CD31 (Miltenyi Biotec, 130-092-652) and CD144 (Miltenyi Biotec, 130-100-742) expression was performed. In most cases, >90% CD31+ were observed. Cultures with lower differentiation efficiency were subjected to CD31+ cells magnetic enrichment using equipment described above and CD31 Microbeads (Miltenyi Biotec, 130-091-935). CD31+ endothelial cells were stored as pellet at -80°C.

##### Trilineage Differentiation

70-80% confluent iPSCs were seeded and differentiated into Ectoderm, Endoderm and Mesoderm using the StemMACS™ Trilineage Differentiation Kit (Miltenyi Biotec), following manufacturer´s instructions. At day 7 of differentiation cells were harvested using Accutase or TrypLE (both Gibco). FACS analysis was performed to characterise ectodermal (SOX2, PAX6), endodermal (CXCR4, SOX17) and mesodermal (CD144, CD140b -Miltenyi Biotec, 130-121-052-) cells. The remaining cells were collected as pellet and stored at -80°C.

#### Immunostaining for FACS

150,000-300,000 single cells were used per tube. Surface marker staining was performed in FACS buffer: DPBS without Calcium and Magnesium (Gibco), BSA 0.5% (Miltenyi Biotec) and 2mM EDTA (Gibco) for 10 minutes at 4°C. After 2 rounds of washing cells were fixed and permeabilized using FoxP3 Staining Buffer Set (Miltenyi Biotec), incubations were done for 30 minutes at 4°C. After washes using Perm/wash buffer cells were incubated with antibodies against intracellular antigens for 60 minutes at 4°C. After final washes cells were resuspended in 250µl FACS buffer. Flow cytometry acquisition was performed using MACSQuant VYB (Miltenyi Biotec) and analysis using FCS Express 5 Plus software.

#### DNA Extraction

DNA extraction was performed on harvested cell pellets using the Maxwell® RSC Instrument (Promega) with either the Maxwell® RSC Cultured Cells DNA Kit (Cat# AS1620) or the Maxwell® RSC Blood DNA Kit (Cat# AS1400) for fresh and frozen cell samples, and the

Maxwell® RSC FFPE Plus DNA Purification Kit (Cat# AS1720) for paraformaldehyde (PFA)-fixed cell samples, with no difference in DNA methylation analysis quality observed. Extracted DNA quantities were measured using the Quantus system together with the "QuantiFluor® ONE dsDNA" kit (Promega).

#### DNA Methylation Analysis

A cohort of n=155 iPSC derivates of eight differentiation states was generated in technical (independent experiments) or biological (distinct cell line donors) replicates, then randomly split into a training cohort of n=97 with a minimum of 8 samples per differentiation state, and a validation cohort of n=58. DNA extraction was performed on each sample cell pellet for subsequent DNA methylation profiling.

#### Array-Based DNA Methylation Profiling

Genomic DNA was bisulfite-converted using the EpiTect Fast Bisulfite Kit (Qiagen) following the manufacturers' protocol. The bisulfite-converted DNA was subsequently analysed using the Infinium MethylationEPIC BeadChip array (v1.0 or v2.0) and scanned on the iScan System (Illumina). Raw DNA methylation data were pre-processed using R (v 4.3.3). Signal intensities were obtained from IDAT files using the Bioconductor package minfi (v 1.40.0) (Aryee et al., 2014). Dye bias correction was performed using the Noob (normal-exponential out-of-band) method with the preprocessNoob function without "between-array normalisation" (Triche et al., 2013). Data filtering steps included the removal of probes with detection p-values greater than 0.05 in more than 10% of samples, probes containing single nucleotide polymorphisms (SNPs) at the CpG site, probes located on sex chromosomes, and cross-reactive probes as previously described (Capper et al., 2018; Chen et al., 2013). Beta values for DNA methylation were calculated from normalised data using the minfi getBeta function.

Copy-number variation analysis from unfiltered methylation array data was performed using the conumee 2.0 Bioconductor package v.2.1.2 (Daenekas et al., 2024). A cohort of 50 control samples displaying a balanced copy-number profile from female donors were used for normalization, to allow for inference of sex from the copy number of the X chromosome.

#### Analysis of Publicly Available Datasets

A publicly available methylation array dataset was systematically curated and downloaded from the following Gene Expression Omnibus (GEO) records as raw signal intensity files (IDAT files):

GSE105093, GSE110544, GSE141521, GSE159479, GSE163322, GSE163324, GSE180402, GSE197723, GSE207076, GSE207108, GSE236202, GSE247428, GSE253190, GSE61461, GSE76372, GSE85828, GSE196339. The PRISMA (Preferred Reporting Items for Systematic Reviews and Meta-Analyses; (Page et al., 2021)) flow chart illustrating the number of datasets identified, included and excluded, and the reason for exclusions is available in Supplementary Figure 7. All samples used in this study are outlined in Supplementary Table 5. GenomicMethylSet objects were generated from raw signal intensity files using the readGEORawFile function of the GEOquery package (v2.62.1). (Davis & Meltzer, 2007). Raw data from EPIC arrays were merged with publicly available 450K array datasets using the combineArrays function in minfi. The combined dataset underwent the same filtering and normalisation process as described above.

#### Differential Methylation Analysis

Differential methylation analysis on the CpG site and region level was performed in a one-to-one fashion between iPSCs and the respective differentiation states as previously described (Maksimovic et al., 2017). Differentially methylated positions (DMPs) were visualised as volcano plots using the package EnhancedVolcano (v 1.20.0). Methylation-specific gene ontology (GO) analysis of promoter-associated differentially methylated positions (DMPs) was conducted for each differentiation state using the gometh function within the missMethyl package (v 1.28.0) (Phipson et al., 2016). Investigation beyond the scope of the classifier i.e. the "broader profiles" was done by identifying the top 20,000 CpGs from our DMP analysis that distinguished each reference differentiation state from iPSC, for iPSC it was the top 20,000 CpGs that were common to all differentiation state vs iPSC comparisons. The annotated tables for the top 20,000 CpGs per differentiation state are available in Supplementary Table 4, full DMP analysis results tables are available on request.

Temporal DNA methylation modules were identified independently for astrocyte and lung differentiation using the same analytical workflow. For each lineage, CpG methylation M-values were analysed using linear models implemented in the R package limma (v. 3.62.2), with differentiation time point included as a categorical variable and parental iPSC line included as a covariate. CpGs exhibiting significant temporal variation were identified using an omnibus moderated F-test, with Benjamini-Hochberg-adjusted $P < 0.05$. To retain biologically meaningful changes, CpGs were additionally required to exhibit a difference of at least 0.10 between the maximum and minimum mean beta values across differentiation stages. For each retained CpG, mean beta values were calculated across samples at each time point and standardized to Z-scores across stages. The resulting temporal methylation trajectories were grouped into two modules using k-means clustering (100 random starts), distinguishing CpGs with decreasing (M1) or increasing (M2) methylation during differentiation. CpGs were annotated using the corresponding Illumina array annotation. Full methylation modules tables are available on request.

#### Unsupervised Analysis

Principal component analysis (PCA) was performed on DNA methylation beta values. Separate analyses were conducted for 4 time points throughout astrocyte differentiation across three parental iPSC lines (n = 12), 6 time points throughout lung differentiation across two parental iPSC lines (n = 12), and an additional 9 time points throughout one astrocyte differentiation with cohorts of reference iPSC and astrocyte samples (n = 35). For each analysis, CpGs with missing or non-finite values were excluded, and the 5,000 most variable CpGs were selected based on their standard deviation across samples. PCA was performed using the prcomp() function in R, with samples as observations and CpGs as variables. The proportion of variance explained by each principal component was calculated from its squared standard deviation relative to the sum across all components. Samples were visualised using PC1 and PC2 with colours indicating differentiation stages. Consecutive stage centroids were connected chronologically to illustrate differentiation trajectories.

Uniform Manifold Approximation and Projection (UMAP) was performed on the top 50,000 most variable CpGs for each respective dataset using the R package umap (v 0.2.10.0). UMAP parameters were set to n_neighbors=18, min_dist=0.3 n_components=2, with a Euclidean distance metric and a fixed random state for reproducibility. The resulting UMAP layout was integrated with sample metadata for downstream analysis. A scatter plot of UMAP coordinates was generated using ggplot2, with points representing individual samples, the final plots depicting the distribution of samples and cell types based on the top 50,000 most variable probes across samples.

#### Classifier development

A random forest classifier was built with ranger (v 0.16.0) within the tidymodels ecosystem (tidymodels v 1.2.0; parsnip v 1.2.1; recipes v 1.0.10; workflows v 1.1.4; tune v 1.2.1; rsample v 1.2.1; yardstick v 1.3.0) under R 4.3.3. Parallel execution used BiocParallel (v 1.36.1) and doParallel (v 1.0-17).

A nested cross-validation framework was used to obtain unbiased estimates of model performance while preventing information leakage (5-fold outer cross-validation repeated three times; 3-fold inner cross-validation).

**Nested cross validation, feature selection and hyperparameter tuning.**

Within each outer training split, CpG sites were first variance-filtered using the training data only, retaining the top 50,000 CpGs ranked by standard deviation to reduce dimensionality. Random forest hyperparameters were tuned exclusively in the inner cross-validation using a 20-point max-entropy (Latin hypercube) design over minimum node size (2-30) and number of trees (300-2000), while mtry was fixed per split to the square root of the number of retained predictors. Hyperparameters were selected by minimising the multiclass Brier score. Using the selected hyperparameters, the model was refit on the full outer training split and permutation importance was computed for all variance-filtered CpGs. CpGs were ranked by permutation importance, and nested feature selection was conducted by fitting models on increasing prefixes of this ranked list (2,000, 5,000, and 10,000 CpGs) using the same inner cross-validation structure. The final feature count per outer split was determined using a one-standard-error plateau rule on the inner-CV Brier score, selecting the smallest feature set whose performance was within one standard error of the minimum. This conservative criterion reduces the likelihood of selecting an overfit model driven by sampling variability.

Across outer splits, selected CpGs were aggregated by selection frequency. The final deployed feature set comprised the top-ranked CpGs up to the median selected feature count across splits. Consensus hyperparameters derived from the nested procedure were then used to refit the model on all training data.

**Probability score calibration and threshold identification.**

For probability calibration, outer fold predictions were pooled, and a multinomial ridge regression model was trained in a cross-fitted manner: predictions for each outer fold were calibrated using a model trained only on predictions from the remaining outer folds. This ensured that no sample was used to fit either the base classifier or the calibrator responsible for its calibrated probabilities. Calibration performance was summarised using the multiclass Brier score.

To determine a rejection threshold, the macro-averaged Youden index was evaluated across candidate probability cutoffs using the pooled cross-fitted calibrated predictions. Uncertainty was quantified by nonparametric bootstrap resampling. The selected operating point was defined as the largest threshold on the macro-Youden plateau, that is, the highest cutoff before any further increase resulted in a statistically significant decrease in macro-Youden based on the 95 percent bootstrap confidence interval. Samples with maximum predicted class probability below this threshold were assigned to a "Not Classifiable" category.

**Final model and variable importance**

The final random forest model was trained on the entire training cohort using the final feature set (10,000 CpGs) and consensus hyperparameters (mtry= 100, min_n = 5, trees = 1192) with a calibrated probability score threshold of 0.5. Variable importance was extracted from the fitted ranger object using the impurity-based measure. The trained classifier was serialised and saved as a workflow object for downstream application.

**Model evaluation**

Performance was assessed using two independent datasets:

1. Internal validation cohort (n = 58).
2. External validation cohort (n = 249, publicly available methylation array data).

The pre-trained random forest classifier was loaded from the serialised workflow object and the underlying ranger model was extracted. The pre-processing recipe was reconstructed using the recipes package and applied to generate a numeric feature matrix and factor response vector. Feature-level contributions for each test sample and a training cohort bias vector were computed from the tidy forest object. Samples from the validation cohorts which received a classification discordant with their annotation were identified. For each such sample, the top CpG features with the most negative contribution toward the annotated class and the most positive contribution towards the SteMClass classification (i.e., features driving their discordant classification) were extracted for further analysis.

**Cumulative-contribution analysis of CpG features**

The tidy representation of the fitted forest (tidyRF) was queried with the featureContrib function from the package tree.interpreter (v 0.1.1), producing a three-dimensional array P × D × N (P = 10 000 CpGs, D = 8 classes, N = number of test samples). For each CpG × class cell we calculated the mean absolute contribution across the N samples, yielding a P × D matrix of average importances. For every class separately, CpGs were then ranked in descending order of this value, and the running sum was divided by the class-specific total to obtain a cumulative-fraction vector. The resulting cumulative curves were combined into a long-format data frame and plotted with ggplot2 (4.0.3) on a log10 scale CpG-rank axis. An 80 % horizontal reference line highlights how the contributions accumulate.

**SteMClass Web Application**

SteMClass was developed in R using the Shiny framework with shiny, shinyjs, shinycssloaders, bslib, and thematic, employing a Bootstrap theme and custom CSS for interface rendering. DNA methylation preprocessing relies on minfi, methylumi, wateRmelon and preprocessCore for raw IDAT file handling and normalisation, with array manifests and annotation (IlluminaMethylationManifest) loaded dynamically. Preprocessing includes noob normalisation, conversion to a common 450K format where required, and genome mapping. The pre-trained SteMClass random forest model (loaded from GitHub Releases (https://github.com/pereze5/SteMClass_App/releases) is applied after aligning test data to the training reference cohort and imputing missing CpGs using means learned during model development. Predictions are output as class probabilities across eight reference cell states, with a rejection option for samples not reaching the 0.5 probability threshold. Visualisations include probability bar charts, UMAP projections generated with uwot, and heatmaps generated with ComplexHeatmap and circlize. Global heatmaps compare the test sample to reference profiles using the top 10,000 CpGs distinguishing a selected differentiation state from iPSC, while gene-level heatmaps display CpGs annotated to a specified gene (UCSC_RefGene_Name) ordered by genomic position and annotated by gene region (UCSC_RefGene_Group). Results and figures are exportable as high-resolution images or HTML outputs.

**Cumulative-contribution analysis of CpG features**

The tidy representation of the fitted forest (tidyRF) was queried with the featureContrib function from the package tree.interpreter (v 0.1.1), producing a three-dimensional array P × D × N (P = 10 000 CpGs, D = 8 classes, N = number of test samples). For each CpG × class cell we calculated the mean absolute contribution across the N samples, yielding a P × D matrix of average importances. For every class separately, CpGs were then ranked in descending order of this value, and the running sum was divided by the class-specific total to obtain a cumulative-fraction vector. The resulting cumulative curves were combined into a long-format data frame and plotted with ggplot2 (4.0.3) on a log10 scale CpG-rank axis. An 80 % horizontal reference line highlights how the contributions accumulate.

#### Nanopore Sequencing

Libraries were prepared with the Rapid Barcoding Kit (Oxford Nanopore Technologies, UK) according to the manufacturer's instructions, using SQK-RBK004 for the 13 samples with paired EPIC profiles and SQK-RBK114.24 for the internal and external validation cohorts. During library preparation, input DNA is fragmented while barcodes are simultaneously attached, using a time-efficient transposase-based approach known as "tagmentation". The final libraries were loaded onto R9.4.1 (FLO-MIN106) flow cells for the EPIC data paired samples and the newer R10.4.1 (FLO-MIN114) flow cells for the validation cohorts. Low coverage whole genome sequencing (WGS) was performed multiplexing up to 6 samples for 6-65 hrs on a MinION Mk 1B device (Oxford Nanopore Technologies, UK). POD5 files containing the raw data were obtained in real time using the manufacturer's software MinKNOW (v.1.3.1-v.3.6.0). Analysis was performed using the nanoDx classification pipeline (v0.6.2) (Kuschel et al., 2022). The SteMClass training cohort raw IDAT files were processed and converted into an H5 file format for use as the training cohort for the nanoDx pipeline using the R package rhdf5 (v2.50.2). The nanoDx classification and analysis pipeline is publicly available at https://gitlab.com/pesk/nanoDx. The source code for the ad hoc random forest implementation is available at https://gitlab.com/pesk/nanoBenchmark. Copy number profiling from nanopore sequencing data was performed using the R package ichorCNA (v.0.5.1) (Adalsteinsson et al., 2017).

#### Nanopore sequencing data processing

POD5/FAST5 files containing the raw data were obtained in real time using the manufacturer's software and raw signal was basecalled with dorado v7.13.6 in MinKNOW v26.01.15 which was set to with the high accuracy 5mCG_5hmCG modified-base model. Reads were aligned to hg19 with minimap2 v2.26. Per-CpG methylation was summarised with modkit pileup v0.2.3 combining calls from both strands of each CpG. 5mC and 5hmC calls were combined into a single modified state, matching the bisulfite-based arrays used to build the reference, which do not distinguish the two. Read-level calls were filtered using modkit's default per-sample threshold estimation, which discards calls below the 10th percentile of call confidence estimated from approximately 10,000 sampled reads. Primary read counts were obtained from the aligned BAM files with samtools v1.17.

#### Classification

Samples were classified with the nanoDx pipeline v v0.6.2; Euskirchen et al., 2017) using an ad hoc random forest trained on the SteMClass reference set (97 samples, 8 classes, 10,000 CpGs). For each sample, reference beta values were restricted to the CpGs covered by that sample and binarised at 0.6. A random forest (scikit-learn 1.3.0: 2,000 trees, entropy criterion, minimum split size 4, random state 42) was trained and wrapped in sigmoid calibration fitted by stratified 5-fold cross-validation. The predicted class and its calibrated probability were recorded. The confidence threshold of 0.5 was prespecified from the microarray SteMClass classifier, samples scoring below 0.5 were designated Not Classifiable. Classification was compared with lineage annotations established by current gold-standard markers.

#### Classification stability analysis.

To determine how much sequencing data a stable SteMClass classification requires, each sequencing run from the validation cohorts which resulted in a classification score meeting the 0.5 cut off was reanalysed retrospectively with increasing amounts of sequencing data (n =63). Each read was assigned its acquisition time from the run sequencing summary file, and the reads were grouped into cumulative subsets at up to 31 predefined time points between 2 and 3,946 min, depending on the length of the run. Time points without CpG features overlapping the reference set were excluded. Stability was determined from 1531 classifications of the 63 included samples, with a median of 24 time points per sample (range, 18-31). The calls were summed to give a cumulative methylation profile at each time point, and each profile was classified with the standard SteMClass workflow. For each time point we recorded the number of reads, the number of CpG features overlapping the reference set, the predicted class and the calibrated score. A classification was considered stable from the first time point after which three conditions held until the end of the run: the predicted class matched the reference class (determined by current gold standard markers), the calibrated score was ≥0.5, and the score stayed within ±0.10 of the sample's end-of-run plateau. The plateau was the median score over all time points with at least 75% of the sample's final number of features. We report the number of features and reads at which 50% and 90% of included samples had a stable classification.
